# Wbm0152, an outer membrane lipoprotein of the *Wolbachia* endosymbiont of *Brugia malayi*, inhibits yeast ESCRT complex activity

**DOI:** 10.1101/2025.07.21.665852

**Authors:** Lindsay Berardi, Alora Colvin, Matthew West, Greg Odorizzi, Vincent J. Starai

## Abstract

Human pathogenic filarial nematodes of the family Onchocercidae, including *Brugia malayi* and *Onchocerca volvulus,* cause debilitating filarial diseases such as lymphatic filariasis and river blindness. These mosquito-borne pathogens are obligately colonized by the gram-negative intracellular alphaproteobacterium, *Wolbachia pipientis,* which is essential for nematode sexual reproduction, long-term survival, and pathogenicity in the mammalian host. Like many intracellular bacteria, *Wolbachia* likely uses numerous surface-exposed and secreted effector proteins to regulate its ability to persist and replicate within nematode host cells. However, due to the inability to cultivate *Wolbachia* in the laboratory and the genetic intractability of both filarial nematodes and the bacterium, the molecular underpinnings that define the bacterium:nematode relationship are almost completely unknown. In this work, we show that the expression of a *Wolbachia* outer membrane lipoprotein, *w*Bm0152, in *Saccharomyces cerevisiae* inhibits the activity of the conserved Endosomal Sorting Complex Required for Transport (ESCRT) complex and strongly disrupts endosomal maturation, leading to defects in ubiquitylated protein turnover. Using in vivo bimolecular fluorescence complementation, we find that Wbm0152 interacts with the Vps2p subunit of the ESCRT-III subcomplex as well as the Vps2p ortholog (BmVps2, Bm6583b) from a *Wolbachia* host nematode, *Brugia malayi*. These data suggest a novel role of ESCRT in *Wolbachia* persistence providing insight into the elusive relationship between these two organisms.

**AUTHOR SUMMARY:** Filarial diseases of mammals, including lymphatic filariasis and canine heartworm, are caused by vector-borne filarial nematodes of the family Onchocercidae. Many of the nematodes in this family are obligately colonized by an intracellular bacterium, *Wolbachia pipientis*, which is essential for the nematode’s long-term survival, reproduction, and pathogenicity. Therefore, understanding the mechanisms used by *Wolbachia to* persist and replicate within host cells could provide new molecular targets for treating filarial infections. Due to the genetic intractability of both nematode and bacterium, however, significant progress on characterizing these interactions have proven difficult. In this work, we show that a predicted outer membrane lipoprotein, Wbm0152, of the *Wolbachia* endosymbiont of *Brugia malayi* inhibits yeast Endosomal Sorting Complex Required for Transport (ESCRT) complex activity in vivo. Wbm0152 interacts with a core subunit of the yeast ESCRT-III complex, as well as with the orthologous ESCRT-III protein from *Brugia*. ESCRTs are conserved across eukaryotes and are important for diverse cellular processes such as endosomal maturation, autophagy, and cellular division. As *Wolbachia* persists within a membrane-bound compartment within *Brugia* and must avoid host autophagic pathways, this study presents a potential mechanism by which *Wolbachia* may regulate *Brugia* membrane trafficking pathways to ensure its intracellular survival.

## INTRODUCTION

Pathogenic filarial nematodes, such as *Brugia malayi, Wuchereria bancrofti*, and *Dirofilaria immitis*, are a group of parasitic, mosquito-borne nematodes that are known to cause debilitating and disfiguring illness in millions of humans and animals worldwide. Typically, humans infected with such nematodes are treated with a regimen of anthelmintic drugs such as ivermectin [1–3]. However, due to ivermectin’s inability to effectively eradicate adult stage worms in vivo [4, 5] combined with increasing observations of anthelmintic resistance in the nematode population [6–8], the demand for identifying novel drug targets to support the elimination of these nematodes has increased rapidly. Interestingly, filarial nematodes of the family Onchocercidae – which include the genera *Brugia, Wuchereria,* and *Dirofilaria* – are colonized by *Wolbachia pipientis,* a gram-negative, obligately intracellular alphaproteobacterium, which is essential for proper nematode reproduction and survival [9, 10]. Elimination of this bacterium from the nematode using doxycycline or tetracycline treatments leads to the sterilization of adult worms, killing of microfilaria, and suppression of infection symptoms in humans [11–14]. Despite the well documented requirement of *Wolbachia* for filarial nematode survival, little is understood about the molecular underpinnings of the complex bacterial:nematode essential endosymbiosis. This lack of knowledge predominantly stems from the inability to cultivate *Wolbachia* in the laboratory and the poor genetic tractability of both *Wolbachia* and these filarial nematodes. Therefore, our knowledge about the *Wolbachia*:nematode relationship stems largely from high-resolution microscopy techniques and heterologous model systems to hypothesize how this bacterium persists within its host.

*Wolbachia*, like many other intracellular bacterial pathogens, including *Legionella pneumophila, Chlamydia trachomatis,* and the closely related alphaproteobacterium *Anaplasma phagocytophilium*, persists and replicates within a host-derived, vacuolar-like compartment [15, 16]. The composition of these bacteria-laden compartments is distinct from normal host organelles and they do not fuse with lysosomes, thereby protecting the bacterium from host degradative pathways. Previous research on both nematode-derived and *Drosophila*-derived *Wolbachia* has suggested that this *Wolbachia*-containing compartment is likely formed from ER or Golgi membranes [17, 18], although the precise lipid and protein composition of this compartment remains unknown. Many intracellular bacteria employ dedicated secretion systems to deploy numerous secreted and surface-exposed proteins – termed effectors – that actively modulate host membrane trafficking and lipid synthesis pathways to create these unique compartments and to inhibit phagosome:lysosome fusion to prevent bacterial degradation [15]. As *Wolbachia* contains a Type IV secretion system [19–22] and survives within an intracellular membrane-bound compartment, it is likely that *Wolbachia* also uses secreted effectors to modulate host membrane dynamics and to ensure the creation of its intracellular replicative niche.

Previously, our laboratory used the unicellular eukaryote *Saccharomyces cerevisiae* as a model system to screen several predicted Type IV-secreted proteins from the *Wolbachia* endosymbiont of *Brugia malayi* for the ability of these candidate effectors to manipulate conserved eukaryotic processes [19, 23]. Expression of one such protein, *w*Bm0152, lead to the aberrant accumulation of an enlarged prevacuolar compartment and the failure to deliver representative endosomal membrane-bound cargo proteins (CPS or Sna3) to the lumen of the degradative vacuole [19]; these phenotypes are similar to those observed in yeast strains defective in <u>e</u>ndosomal <u>s</u>orting <u>c</u>omplexes <u>r</u>equired for transport (ESCRT) activity [24]. ESCRTs are highly conserved, multisubunit complexes that are responsible for the invagination and scissioning of cellular membranes *away* from the cytosol. In yeast, ESCRTs are essential for intralumenal vesicle formation (ILV) during endosome maturation and for microautophagy [25–28]. In mammalian cells, ESCRTs are important for these same functions as well as membrane repair [29], nuclear envelope remodeling [30, 31], and viral budding [32–34].

ESCRT subcomplexes assemble in an ordered, stepwise fashion in which the first three complexes (ESCRT-0, -I, -II) function to bind and cluster ubiquitylated proteins on endosomal membranes, as well as recruiting and binding protein subunits of the downstream ESCRT complex. The fourth and most highly conserved complex (ESCRT-III) has the unique job of inducing membrane deformation and scissioning to allow for the creation of the endosomal ILVs. ESCRT-III assembly on endosomes is initiated when myristolated Vps20p is recruited to the endosomal membrane by the assembled ESCRT-II complex (consisting of Vps22p, Vps25p, and Vps36p), which then recruits and initiates the homopolymerization of Snf7p. The Vps2p:Vps24p module of ESCRT-III binds to the Snf7p homopolymer, extending its polymerization laterally, thus creating a three-dimensional helical structure that induces membrane curvature towards the lumen of the endosome [35–40]. The ESCRT-III accessory protein, Bro1p, also binds directly to Snf7p, promoting the recruitment of the ubiquitin hydrolase, Doa4p, which removes and recycles the ubiquitin moiety from target proteins prior to ILV internalization. Finally, the AAA+ ATPase, Vps4p, is recruited to disassemble the ESCRT complexes and completes the scission of the ILV vesicle into the lumen of the endosome [41–43]. Disruption of most of the core ESCRT proteins results in the aberrant accumulation of flattened stacks of endosomes that contain few ILVs and fail to properly fuse with the vacuole/lysosome; these abnormal endosomes have been termed ‘class E compartments’ [44].

In this study, we now show that *w*Bm0152 expression inhibits the ILV-formation activity of ESCRT in yeast. By using the rapamycin-induced degradation (RapIDeg) system [45] to visualize ESCRT activity in vivo, we show that *w*Bm0152 expression prevents the ESCRT-dependent translocation of an artificially-ubiquitylated vacuole membrane protein into the lumen of the degradative vacuole. We also demonstrate that expression of *w*Bm0152 inhibits the formation of ILVs in yeast endosomes using cryo-fixation tomography. Using bimolecular fluorescence complementation assays, we find that Wbm0152 binds in vivo to the yeast ESCRT-III protein Vps2p – as well as the *B. malayi* Vps2p ortholog – and we show that Wbm0152 alters the recruitment of the downstream ESCRT accessory proteins through colocalization studies and electron microscopy. These findings not only identify a bacterial protein capable of inhibiting ESCRT activity but also illuminate an intricate relationship between *Wolbachia* and its nematode host to support its intracellular survival.

## RESULTS

### Wbm0152 inhibits ESCRT-dependent protein degradation in yeast

Our previous research showed that expression of *w*Bm0152 in yeast inhibited the normal delivery of the endosomal cargo proteins carboxypeptidase S (CPS) and Sna3p to the degradative vacuole lumen; these phenotypes are also observed in ‘class E’ protein sorting mutants of yeast caused by defects in Endosomal Sorting Complex Required for Transport (ESCRT) subunits [19, 46, 47]. Therefore, we hypothesized that *w*Bm0152 expression may be inhibiting ESCRT either directly or indirectly. To test this hypothesis, we utilized the Rapamycin Dependent Degradation Assay (RapIDeg) [45] to determine the impact of *w*Bm0152 expression on the ESCRT-dependent vacuolar degradation of an artificially-ubiquitylated protein. In this assay, cells express the vacuolar membrane iron transporter, Fth1p, are fused to a GFP and FK506 Binding Protein (FKBP) domain. These strains also harbor the FKBP-rapamycin binding domain (FRB) fused to 3x ubiquitin, which allows the dimerization of the FRB and FKBP domains in the presence of rapamycin, causing the artificial ubiquitylation of the Fth1-GFP protein. This ubiquitylation recruits ESCRT complexes to the vacuole membrane, thus concentrating the Fth1-GFP-3xUb cargo and delivering Fth1-GFP-3xUb into the vacuolar lumen in an ESCRT-dependent manner, leading to the degradation of the Fth1-GFP cargo [45, 48].

In empty vector control RapIDeg strains, we observed clear vacuolar membrane localization of Fth1-GFP, as expected (**Fig. 1A**). After rapamycin addition, we observed approximately 90% of the cell population move Fth1-GFP into the vacuole lumen, with concomitant degradation of that protein (**Figs. 1A and B**). In contrast, strains expressing *w*Bm0152, failed to degrade Fth1-GFP with only 2% of cells accumulating Fth1-GFP in the vacuole lumen 2h after treatment with rapamycin (**Fig. 1A and B**), showing that *w*Bm0152 inhibits the ESCRT-dependent invagination of the vacuolar membrane in response to a ubiquitylated membrane protein. Therefore, *w*Bm0152 expression prevents the ESCRT-dependent turnover of a ubiquitylated protein in yeast.

**Figure 1.**
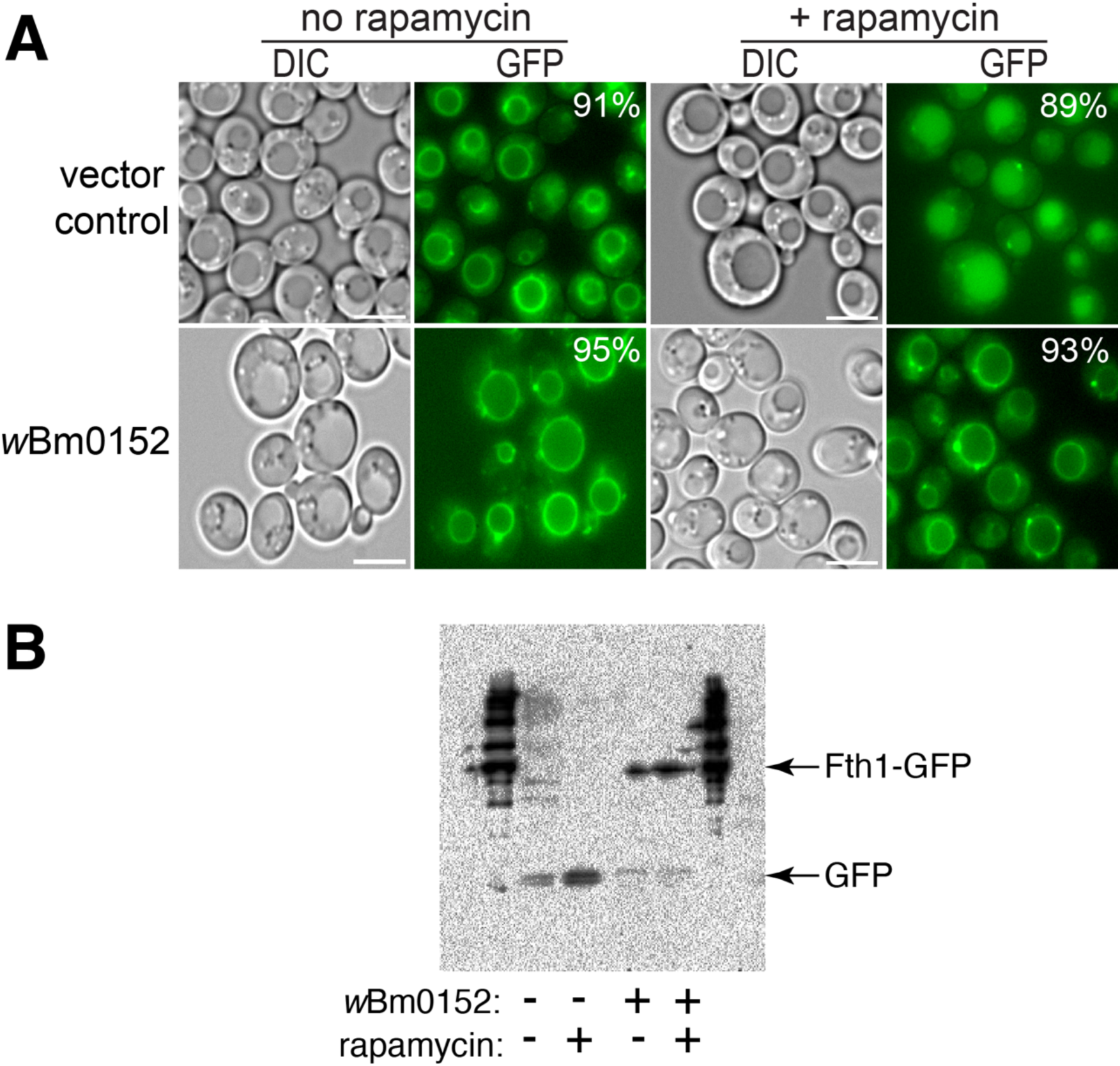
Wbm0152 expression inhibits ESCRT-dependent ubiquitylated protein turnover. RapIDeg yeast strains harboring either a vector control or the β-estradiol inducible *w*Bm0152 expression vector were assayed for vacuolar turnover of Fth1-GFP-FKBPx2, as in Methods. **A)** Representative images showing localization of Fth1-GFP in response to 1 µg mL^−1^ rapamycin; bar = 5 µ. Listed percentages indicate the population of cells displaying the presented Fth1-GFP localization; *n* > 400 cells per condition. **B)** Representative α-GFP immunoblot of whole cell lysates prepared from cells in **A)**.

### Expression of *w*Bm0152 in yeast induces ESCRT mutant-like growth defects

Previously, our lab observed a slight growth defect of *w*Bm0152-expressing yeast strains in the presence of zinc and caffeine, suggesting expression of Wbm0152 may alter endolysosomal membrane compartment function [19]. Based on our RapIDeg results indicating that *w*Bm0152 likely inhibits ESCRT activity, we sought to determine if *w*Bm0152 expression induced specific growth phenotypes like those observed in yeast strains lacking individual ESCRT subunit proteins. Previous research identified that the 1,3-β-glucan-binding azo dye, Congo red [49], inhibited the growth of several individual ESCRT subunit mutants [50]. This same study also revealed that many of the ESCRT-III or ESCRT-III related subunits have distinct levels of growth inhibition when grown on media containing Congo red. Particularly, ESCRT-III mutant strains such as *vps2Δ* and *vps24Δ* are exquisitely sensitive to the presence of Congo red, while other strains – like *snf7Δ* and *vps20Δ* – are only moderately sensitive to these conditions, allowing us to compare growth patterns between ESCRT subunit mutants and *w*Bm0152 expressing strains.

Vector control strains are insensitive to the presence of 15 µg ml^−1^ Congo red on minimal medium (**Fig. 2**). Strains constitutively expressing *w*Bm0152, however, are strongly inhibited for growth under these conditions (**Fig. 2**). When comparing the growth of this strain to representative deletion mutants of ESCRT complex subunits, we noted that our *w*Bm0152-expressing strain has similar levels of sensitivity to Congo red as *vps2Δ*, *vps24Δ,* and *bro1Δ* strains; no other ESCRT subunit deletion strain showed comparable growth sensitivities under these conditions. These results indicate that expression of *w*Bm0152 likely impacts the function of the yeast ESCRT-III subcomplex, as well as the downstream ESCRT accessory proteins (Bro1p) required for efficient ESCRT function. Vps2p and Vps24p are known to interact in a bipartite module, binding to polymerized Snf7 to promote the creation of the 3-dimensional helix to induce endosomal membrane deformation and ILV formation [35]. Bro1p, known to be activated by the polymerization of Snf7, is a key accessory protein required for the recruitment of the ubiquitin hydrolase, Doa4p [51]. In addition to its role in recruiting Doa4p to ESCRT-III, Bro1p:Snf7p interactions also inhibit the association of Vps4p with the ESCRT complex, allowing time for the deubiquitylation of protein cargo by Doa4p [52]. Therefore, Wbm0152 may be disrupting the recruitment of the downstream ESCRT accessory proteins to the assembled complex. As ESCRT-III is primarily responsible for the physical deformation of the endosomal membrane to generate ILVs, we next chose to directly observe the impact of *w*Bm0152 expression on yeast endosomal maturation via electron microscopy.

**Figure 2.**
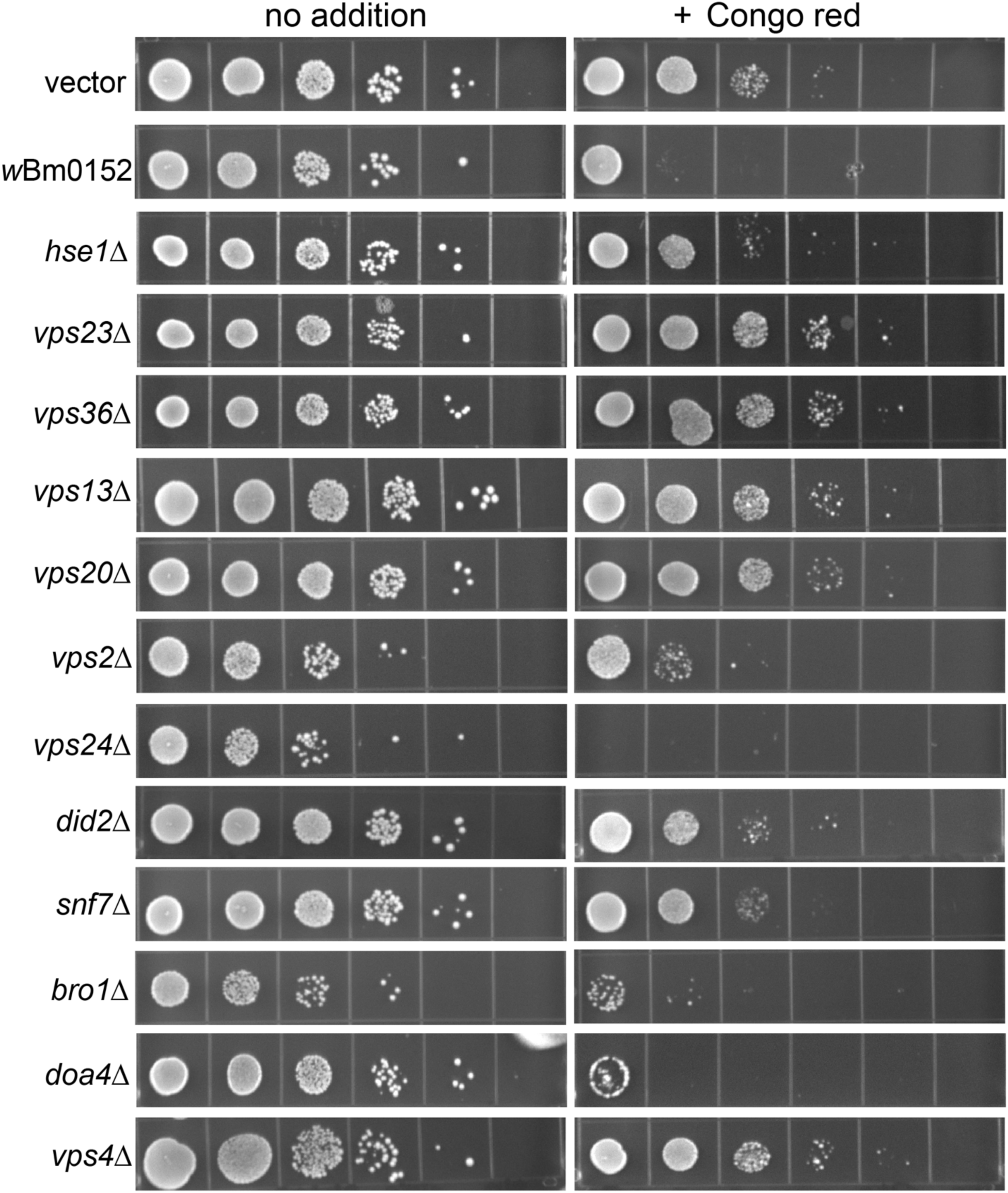
Wbm0152 expression phenocopies ESCRT mutant growth defects. BY4742 yeast strains harboring the indicated ESCRT subunit deletion, an empty vector control, or the constitutively expressing pYES*_TDH3_*-*w*Bm0152 (Methods) were grown overnight at 30 °C in CSM-URA media. Cultures were diluted to a final OD_600_ = 1.0, serially diluted 10-fold four times into sterile water, and 10 µL of each dilution was spotted to CSM media lacking uracil either lacking or containing 15 µg mL^−1^ Congo red. Plates were incubated at 30°C and imaged after 72h.

### wBm0152 expression results in defective ILV formation and endosomal morphology

During endosomal maturation, ESCRT activity is required to create the ILVs indicative of late endosomes (also termed multivesicular bodies, or MVBs). To visualize to impact of *w*Bm0152 expression on endosomal maturation, we used cryo-fixation for electron tomography to generate high-resolution images of endolysosomal membrane compartments and associated organelles. We observed spherical MVBs in both the control strain and the *w*Bm0152-expressing strain, although the strain harboring *w*Bm0152 appeared to have a slight reduction in the numbers of MVBs per 100 cell profiles (**Figs. 3A-C and D-F**, quantified in **Fig. 3G**). Notably, we also found tubular MVBs in strains expressing *w*Bm0152 (**Figs. 3E and 3F**), similar to those found in other ESCRT-impaired yeast strains like *vta1Δ, did2Δ, vps60Δ,* or *BRO1*-overexpressing strains [42, 43, 52–58]. Furthermore, we noted numerous aberrant membrane structures in *w*Bm0152-expressing cells when compared to control strains, including tubular endoplasmic reticulum and increased frequency of lipid droplets (**Figs. 3E and 3F**).

**Figure 3.**
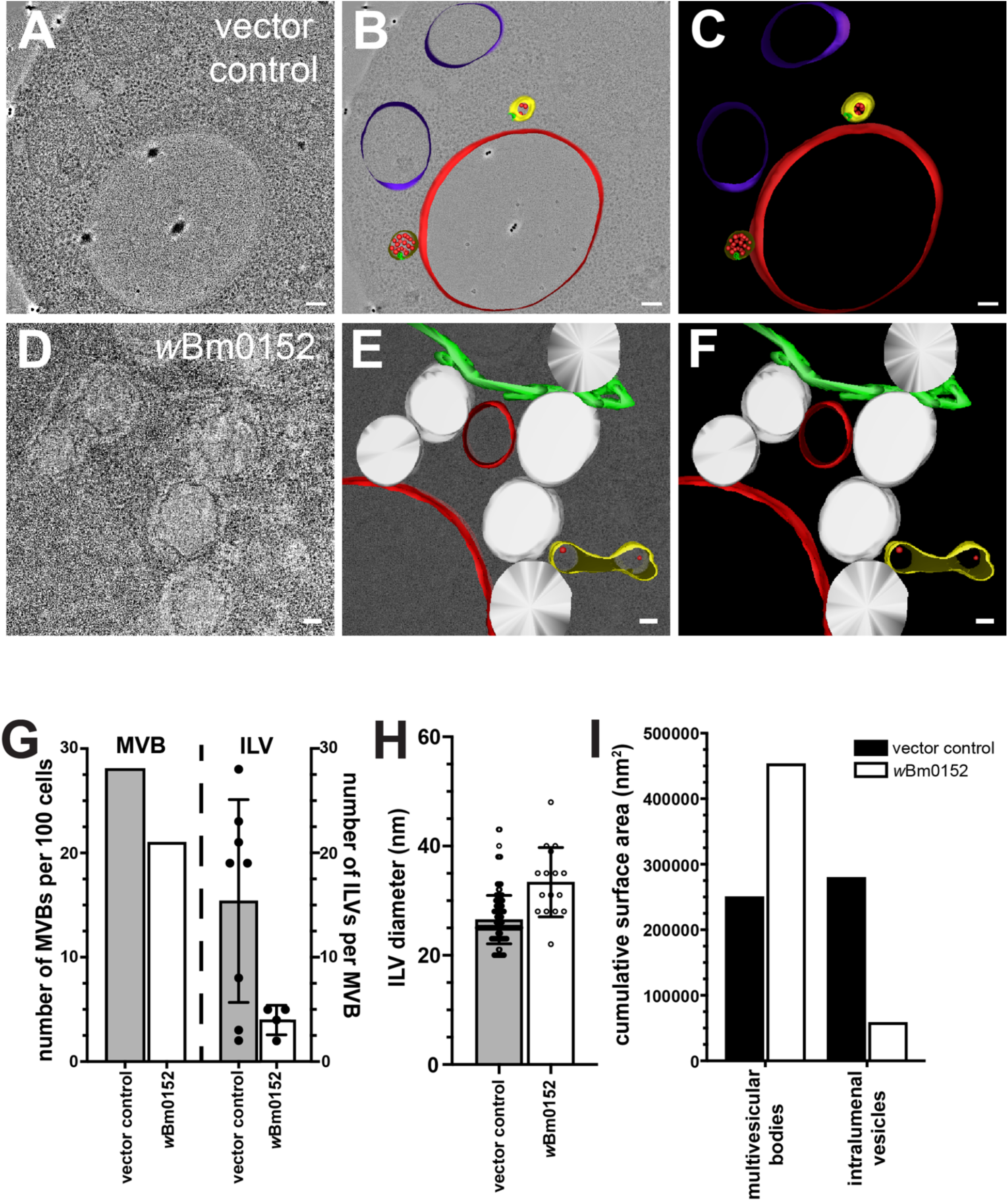
*w*Bm0152 expression inhibits intralumenal vesicle formation in vivo. Yeast strains harboring either the vector control (**A-C**) or *w*Bm0152 expression vector (**D-F**) were visualized via cryo-fixation electron tomography (Methods, bar = 100 nm) and membrane structures were modeled as indicated: MVBs, yellow; degradative vacuole, red; endosomal ILVs, red spheres; ER, green; lipid droplets; white. **G)** Quantification of total MVBs (left) or ILVs per MVB (right) observed across 100 cell profiles for the indicated yeast strain. **H)** Calculated diameters of all ILVs observed in cell profiles from **(G)**. **I)** The total surface area of the limiting membranes of observed MVBs or ILVs from the tomogram profiles of the indicated strains were quantified, as outlined in Methods.

To better observe the morphology of the smaller and less frequent MVBs found in *w*Bm0152-expressing strains, we modeled the membrane compartments found in each tomographic stack and calculated the physical dimensions of the relevant structures. Compared to control strains, the endosomes in strains harboring *w*Bm0152 contained fewer ILVs per MVB (**Fig. 3G**), although those infrequent ILVs tended to be larger in diameter than those found in control strain MVBs (33.38 nm *vs* 26.51 nm, P<0.0001; **Fig. 3H**). By calculating the surface area of the limiting membranes of the MVBs and ILVs in both strains, we found that the cumulative surface area of the MVBs in *w*Bm0152-expressing strains was approximately twice that of the control strains with a concomitant reduction in ILV membrane surface area (**Fig. 3I**). Therefore, *w*Bm0152 expression strongly inhibits the formation of endosomal ILVs in vivo, providing additional evidence of its ability to inhibit ESCRT complex activity.

### Wbm0152 colocalizes with representative ESCRT complex subunits

To determine if Wbm0152 localizes with ESCRT protein subunits in vivo, we performed a colocalization analysis with GFP-tagged candidate ESCRT protein subunits and Wbm0152-mRuby. Each ESCRT-GFP subunit (Vps27-GFP, ESCRT-0; Vps36-GFP, ESCRT-II; Snf7-GFP, ESCRT-III; Bro1-GFP, accessory) was localized to multiple punctate structures indicative of endosomal compartments, as expected [36, 51, 59, 60] (**Fig. 4A**). Analysis of these strains co-expressing *w*Bm0152-mRuby showed strong colocalization of Wbm0152-mRuby with Vps27-GFP, Vps36-GFP, and Snf7-GFP, as measured by Pearson Correlation Coefficient (**Figs. 4A and 3B**). Interestingly, however, Wbm0152 failed to strongly colocalize with the ESCRT accessory protein Bro1-GFP (**Figs. 4A and 4B**). As it is known that Bro1p normally binds to – and colocalizes with – Snf7p to direct the recruitment of Doa4p to the assembled ESCRT complex [51, 52], it was surprising to note that Bro1p did not strongly colocalize with Wbm0152, despite the observation that Snf7p colocalized with Wbm0152. These results show a reduction in the recruitment of Bro1p to assembled ESCRT complexes in the presence of Wbm0152, which may result from Wbm0152-dependent ESCRT-III complex assembly or disassembly defects.

**Figure 4.**
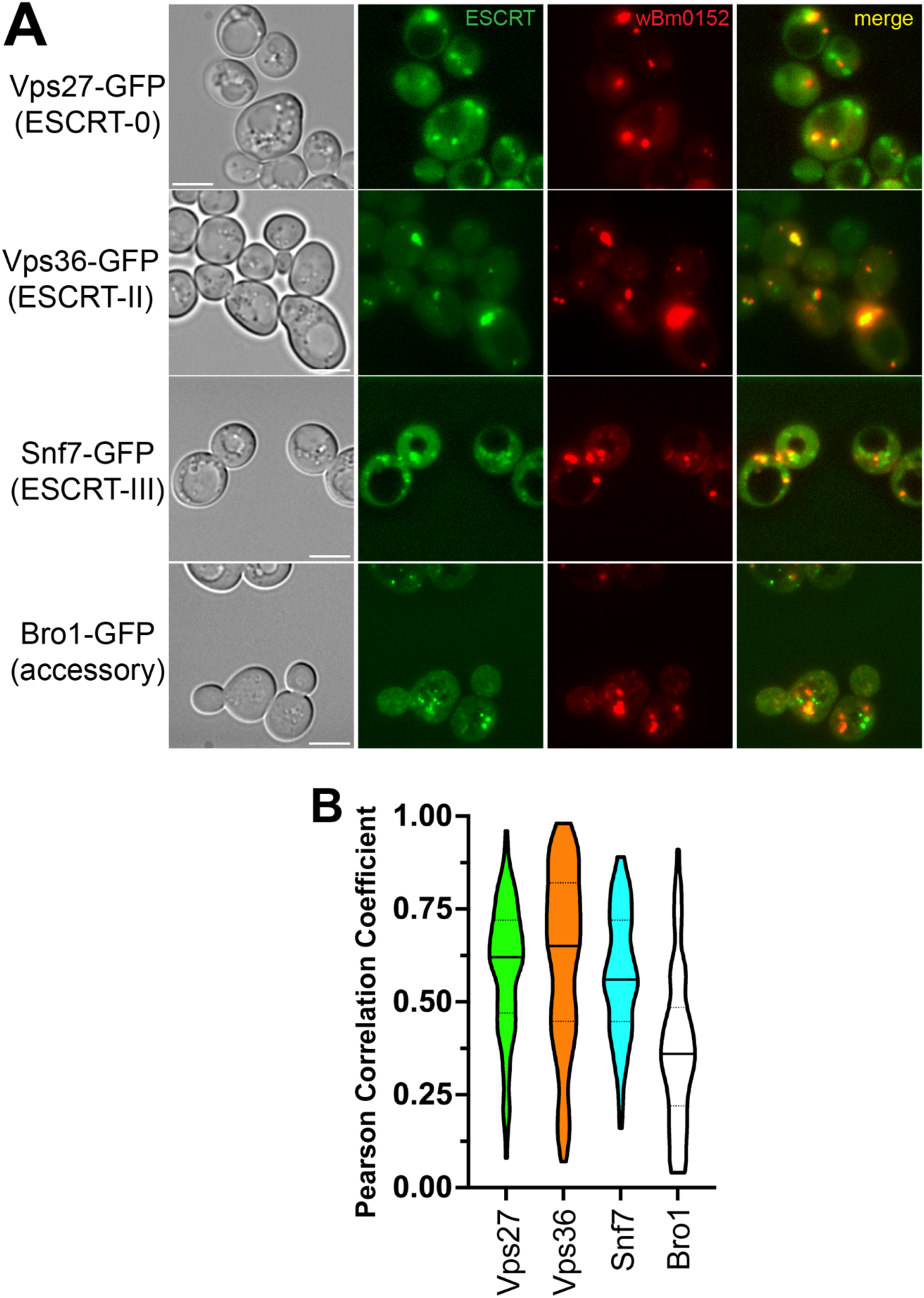
wBm0152 colocalizes with core ESCRT subunits in vivo. **A)** SEY6210 yeast strains harboring the indicated chromosomal ESCRT subunit-GFP and the pYES-*w*Bm0152-mRuby expression vector were grown to saturation at 30°C in selective media. Cells were diluted 1:10 into fresh selective media containing 1 µM β-estradiol, outgrown for 6h at 30°C, and imaged. Bar = 5 µ. **B)** Truncated violin plot showing calculated Pearson’s Correlation Coefficient of the ESCRT:*w*Bm0152 colocalization.

### Wbm0152 interacts with ESCRT-III protein Vps2p

As it appears that *w*Bm0152 expression either inhibits the activity of ESCRT-III or disrupts ESCRT complex assembly dynamics, we hypothesized that Wbm0152 would interact with at least one protein subunit of the ESCRT-III complex. By utilizing a split-Venus bimolecular fluorescence complementation assay (BiFC) [61–63], we sought to identify potential ESCRT binding partners of Wbm0152 through the reconstitution of fluorescence upon protein:protein interactions. Therefore, yeast strains harboring individual ESCRT protein fusions with either the non-fluorescent Venus N-terminal (VN) or Venus C-terminal (VC) domains were transformed with a copper-inducible expression construct containing *w*Bm0152 fused to the complementary Venus fragment. Should Wbm0152 interact with any of these ESCRT subunits and bring the VN and VC domains into close proximity, the fluorescent Venus chromophore will be reconstituted and detectable via fluorescence.

In yeast strains expressing Snf7-VC and Vps2-VN – proteins from the ESCRT-III complex known to directly interact in vivo [42] – we detected numerous fluorescent punctae, as expected (**Fig. 5**); these punctae were not observed in strains harboring the Snf7-VC and empty vector control constructs (**Fig. 5**). Using the inducible *w*Bm0152-VN (or-VC) expression construct from above and a number of ESCRT subunit strains expressing the reciprocal Venus fusion, we identified fluorescent punctae under *w*Bm0152-VC induction conditions only in Vps2-VN harboring strains (**Fig. 5**); no other subunit tested showed fluorescence under similar conditions (**Fig. S1**; tested Vps25p (ESCRT-II), Vps20p (ESCRT-III), Vps24p (ESCRT-III), Snf7p (ESCRT-III), Bro1p (accessory), the Snf7p-binding domain of Bro1p (BOD [42]), and Doa4p). Interestingly, we did not see reconstitution of Venus fluorescence in Vps2-VC/*w*Bm0152-VN strains (**Fig. S1**). This observation could be due to the C-terminal -VN fusion forcing Vps2 into a constitutively active, ‘open’ conformation to bind Wbm0152, which is known to occur with yeast and mammalian ESCRT-III GFP fusion proteins [40, 64, 65]. Alternatively, the VC/VN components in this putative Vps2p-Wbm0152 interaction may simply be in the wrong orientation to form the fluorescent chromophore upon interaction. Regardless, these results show that *Wolbachia* Wbm0152 likely interacts specifically with the Vps2p ESCRT-III subunit in vivo.

**Figure 5.**
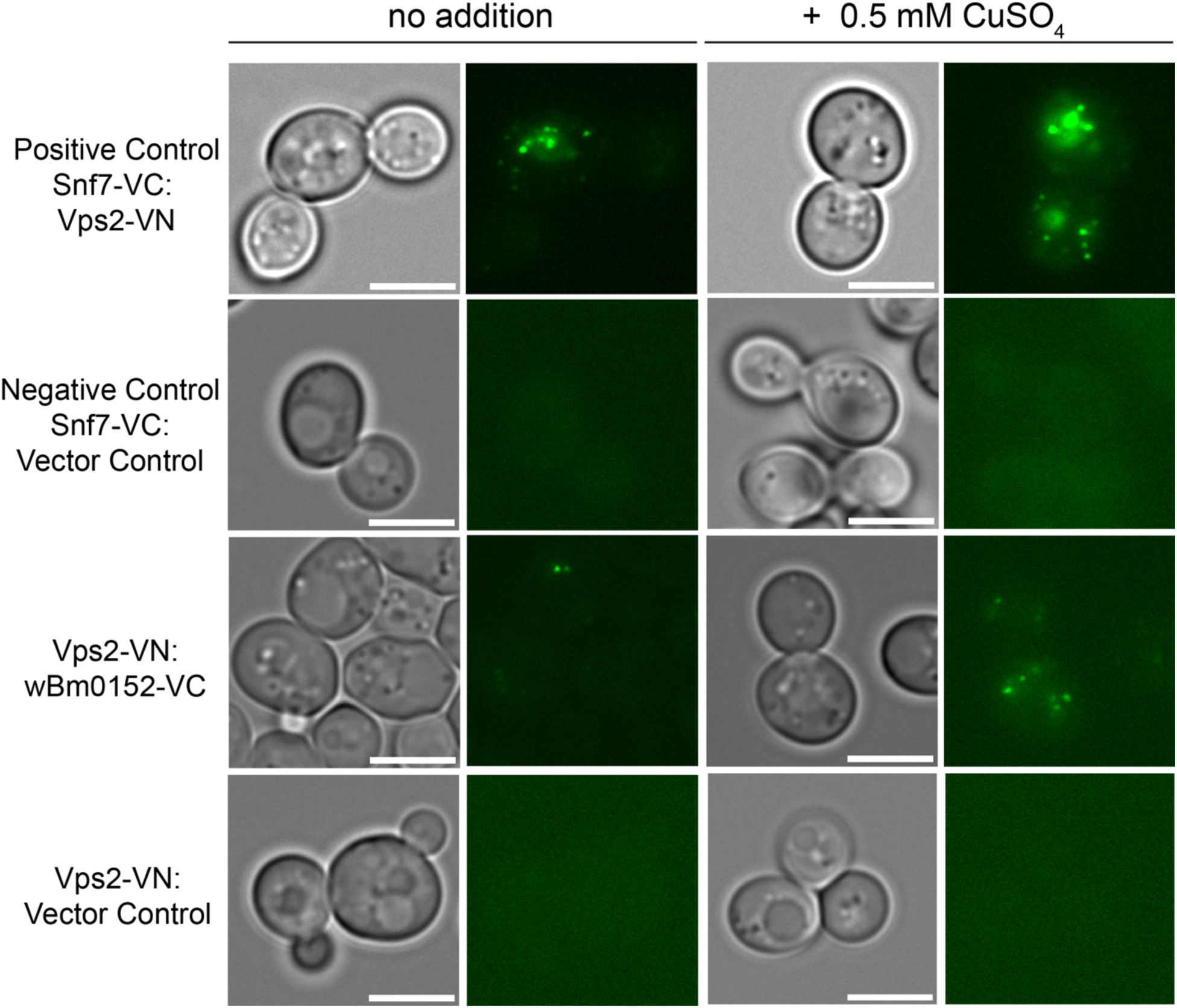
wBm0152 binds ESCRT-III subunit Vps2 in vivo. SEY6210 strains harboring either the N-terminus (VN) or C-terminus (VC) of a Venus-YFP molecule on the C terminus of the indicated ESCRT subunit were transformed with the corresponding copper-inducible pYES*_CUP1_*-*w*Bm0152-VN or pYES*_CUP1_-w*Bm0152-VC plasmid. Strains were grown in CSM media lacking uracil for 18h at 30°C with shaking, diluted 1:10 into fresh selective media lacking or supplemented with 0.5 mM CuSO_4_, and outgrown for 6 hours before imaging. Bar = 5 µ; images are representative of three separate experiments.

### Wbm0152 requires normal ESCRT assembly for activity

Taking advantage of the growth defect we observed on media containing Congo red upon *w*Bm0152 expression in a wildtype strain background, we sought to determine which ESCRT components were required for the Wbm0152-mediated toxicity on this medium. Therefore, candidate ESCRT deletion strains representing one protein from each of the ESCRT-0,-I, and -II complexes – as well as each subunit of the ESCRT-III and downstream accessory complexes – were grown on media containing Congo red with and without *w*Bm0152 expression. As we observed previously, strains missing components of the ESCRT-0, -I, and -II complexes harboring a vector control did not show strong growth defects in the presence of Congo red, when compared to the vector control wild type strain (**Fig. 6**). Strikingly, *w*Bm0152 expression in these mutant backgrounds did not cause additional growth defects on Congo red, in contrast to the wild type (**Fig. 6**). Similarly, *w*Bm0152 expression did not induce Congo red growth defects in *vps20Δ* or *snf7Δ* strains (**Fig. 6**). As each ESCRT subcomplex generally does not assemble and accumulate on the endosomal membrane without the proper assembly and recruitment of the previous ESCRT subcomplex [40, 59, 60, 66, 67], these results show that normal, ordered ESCRT complex assembly is required for *w*Bm0152 activity under these growth conditions. ESCRT disassembly was not important for the toxicity of *w*Bm0152 expression, however, as *vps4Δ* strains remained sensitive to *w*Bm0152 expression on Congo red (**Fig. 6**). Due to the strong growth defect observed on Congo red in *vps2Δ*, *vps24Δ*, *bro1Δ*, and *doa4Δ* strains alone, the necessity of these subunits for Wbm0152 toxicity could not be determined. Taken together, these data suggest the action of Wbm0152 requires fully assembled ESCRT complex and acts upstream of the *vps4Δ* disassembly apparatus.

**Figure 6.**
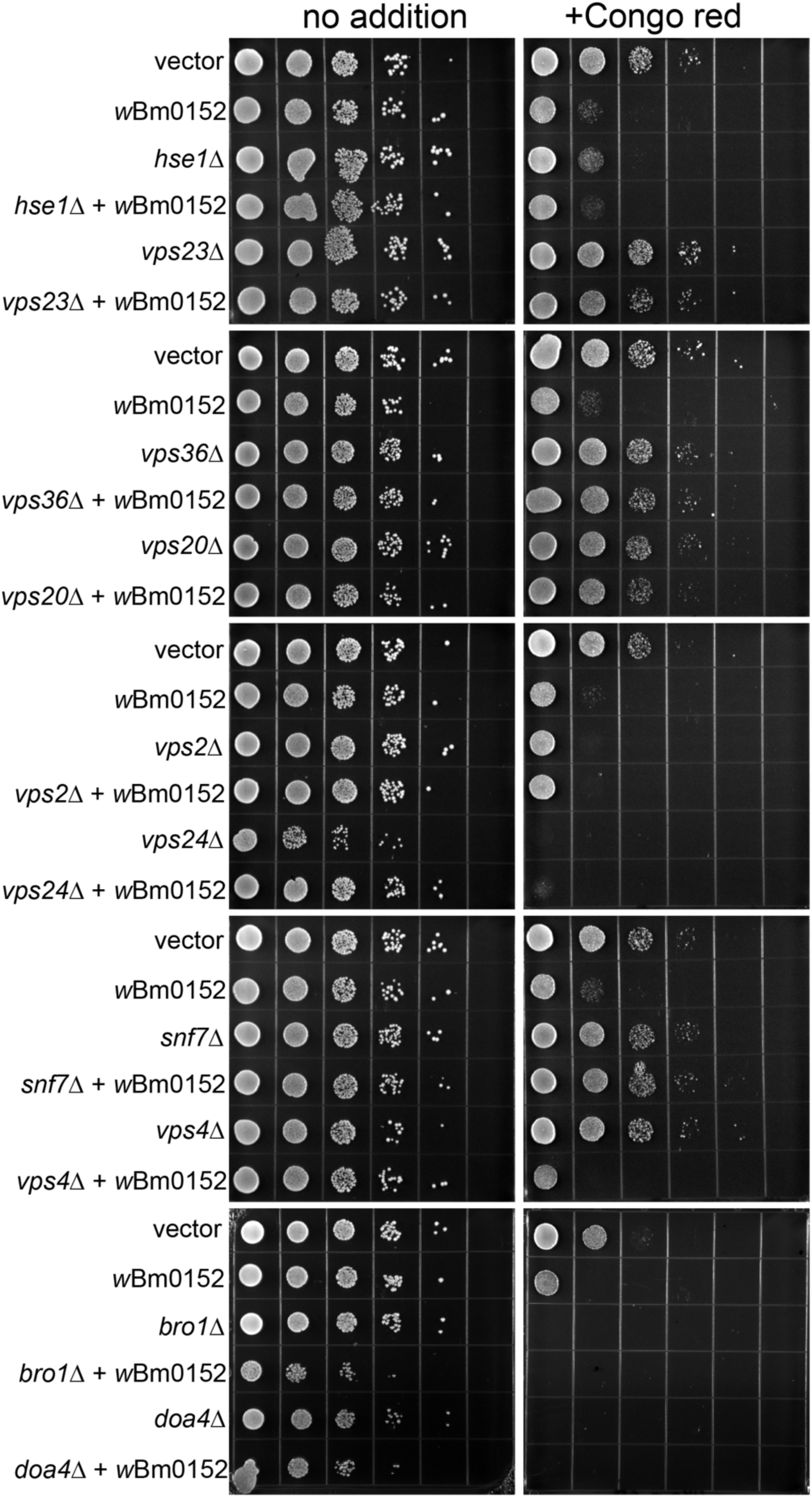
Wbm0152 activity requires ESCRT assembly. BY4742 yeast strains harboring the indicated ESCRT subunit deletion, an empty vector control, or the constitutively expressing pYES*_TDH3_*-*w*Bm0152 (Methods) were grown overnight at 30 °C in CSM-URA media. Cultures were diluted to a final OD_600_ = 1.0, serially diluted 10-fold four times into sterile water, and 10 µL of each dilution was spotted to CSM media lacking uracil either lacking or containing 15 µg mL^−1^ Congo red. Plates were incubated at 30°C and imaged after 72h.

### Wbm0152 interacts with the *Brugia* Vps2 ortholog in yeast

We initiated this work to identify an interaction between a *Wolbachia* candidate effector protein, Wbm0152, with a conserved eukaryotic protein from yeast, which would help elucidate the activity of Wbm0152 in the *Wolbachia*:*Brugia malayi* endosymbiosis. As ESCRT complexes are broadly conserved across eukaryotes, we identified the *VPS2* ortholog from the *B. malayi* genome (*BM6583,* isoform b) [68] and constructed a yeast codon-optimized expression vector to assess Wbm0152:Bm6583b interactions in vivo.

As it is known that Vps2p interacts directly with Snf7p during ESCRT-III complex formation and that interaction can be observed using BiFC, we first ensured that Bm6583b (hereafter, *Bm*Vps2) would interact with yeast Snf7p in vivo. As observed previously, a yeast strain expressing Snf7-VC and Vps2-VN reconstitutes Venus fluorescence (**Fig. 7A**), confirming the expected Vps2p:Snf7p interaction in vivo. Expression of *Bm*Vps2-VN from the endogenous yeast *VPS2* promoter in the Snf7-VC background also produced fluorescent punctae (**Fig. 7A**), showing that *Bm*Vps2 also interacts with yeast Snf7p and likely engages in ESCRT-III complexes with the yeast protein subunits. Next, we introduced the inducible *w*Bm0152-VC vector previously used into strains expressing *Bm*Vps2-VN in both wild type and *vps2Δ* backgrounds. In both backgrounds, we observed the reconstitution of Venus fluorescence only upon induction of *w*Bm0152-VC expression (**Fig. 7B**), showing that *w*Bm0152 interacts with *Bm*Vps2 in a similar manner to yeast Vps2p in vivo, thus strengthening the possibility that Wbm0152 interacts with ESCRT-III subunits in the nematode.

**Figure 7.**
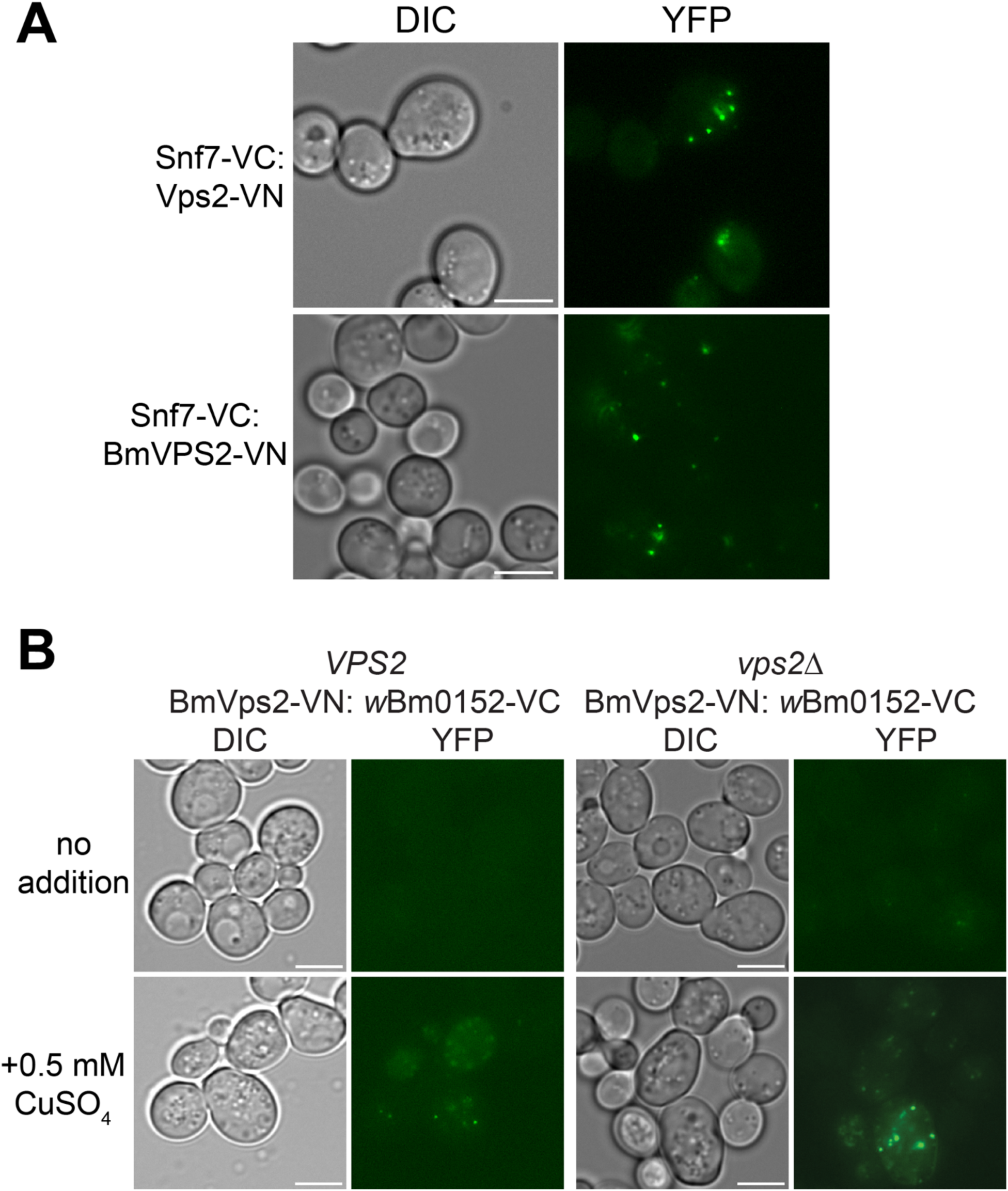
wBm0152 binds *Brugia malayi* Vps2 ortholog (Bm6583b) in yeast. **A)** Yeast SEY6210 harboring yeast wild type SEY6210 (left) or *vps2Δ* (right) strains harboring the plasmids pRS414-pVPS2-*Bm6583b*-VN (Methods) and pYES*_CUP1_-w*Bm0152-VC were grown in CSM media lacking uracil and tryptophan for 18h at 30°C with shaking, diluted 1:10 into fresh selective media lacking or supplemented with 0.5 mM CuSO_4_, and outgrown for 6 hours before imaging. Bar = 5 µ; images are representative of three separate experiments.

## DISCUSSION

In this study, we have shown that expression of Wbm0152 in yeast inhibits the eukaryotic ESCRT complex via interactions with the core ESCRT-III protein, Vps2p. This inhibitory activity requires the formation of a properly assembled ESCRT complex, as in the absence of ESCRT-0, -I, -II, and most -III subunits, *w*Bm0152 expression failed to induce growth defects on media containing Congo red. Importantly, Wbm0152 was also found to interact with the *Brugia* Vps2 ortholog in yeast (Bm6583b), providing evidence of a physiologically-relevant interaction within the nematode that cannot be easily confirmed in the natural host. These data highlight a potential role for Wbm0152 in the regulation of *Brugia* ESCRT activity during endosymbiosis.

Despite the many advances made in the analysis of *Wolbachia*:host interactions, much of the current research focuses on understanding *Wolbachia* survival and transmission in arthropod hosts, in which the relationship between the bacterium and host is usually parasitic in nature. Through a combination of mutant insect lines and cell culture systems, researchers have shown the importance of host actin dynamics for the uptake and maternal transmission of *Wolbachia* during host development [69, 70], although information regarding *Wolbachia’*s ability to <u>actively</u> manipulate these host pathways remains lacking. Understanding of the *Wolbachia*:nematode relationship, however, remains even less clear due to the genetic intractability of *Brugia* and *Wolbachia*, the general difficulty to rear such nematodes, the lack of primary or immortalized nematode cell lines, and the fact that nematode-derived *Wolbachia* (strain types C and D) is known to express evolutionarily-divergent Type IV-secreted effectors than the arthropod-derived *Wolbachia* (strain types A and B) [71]. Therefore, the development of an alternative biological model system for the study of these *Wolbachia* effector proteins found in filarial nematodes is demanded. Our lab, as well as several others, have had great success utilizing *Saccharomyces cerevisiae* as a model eukaryotic cell to study the molecular underpinnings of many host:bacterium interactions, especially when the target of these bacterial effectors is conserved in eukaryotes [72–74].

Many intracellular bacteria such as *Salmonella enterica* and *Mycobacterium tuberculosis* are known to directly modulate host ESCRT activity, likely to support intracellular survival and transmission. Although there are well documented ESCRT-interacting proteins, like the *Mycobacterium* secreted effectors EsxG and EsxH [75], ESCRT-interacting domains amongst intracellular bacteria are not well conserved, nor are their molecular activities on ESCRT well known. Despite the lack of understanding on how intracellular bacteria interact with ESCRT, there is a wealth of knowledge on how many viruses, including HIV-1, interact with and manipulate ESCRT to promote viral budding within mammalian cells. Most of these viral effector proteins are known to contain P(S/T)AP, PPxY, or YPx(n)L motifs that mimic motifs present on ESCRT [76]. The P(S/T)AP domain is known to interact with TSG101, the human ortholog to the yeast ESCRT-I protein Vps23p [77–79]. The PPxY motif is noted to interact with NEDD4, the human ortholog to the yeast Rsp5p protein [80–82], which is a ubiquitin ligase that interacts with Vps27p and Hse1p to regulate the biogenesis of MVBs [83, 84]. Lastly, the YPX(n)L motif is known to interact with ALIX (yeast Bro1p), [64] which will also be found in complexes containing the ESCRT-III subunits CHMP2, (Vps2p ortholog), CHMP4 (Snf7p ortholog), and Vps4p (reviewed in [85]). Some of these domains have been found on ESCRT-interacting proteins from other intracellular pathogens, such as GRA14 and RON4 from *Toxoplasma gondii*, which are known to contain the P(T/S)AP and YPX(n)L domains, respectively [86, 87]. However, it is important to note that *Toxoplasma* has other known ESCRT-interacting proteins, such as GRA64, which is noted to interact with ESCRT subunits TSG101, VPS37, VPS28, UMAD1, ALG-2, and CHMP4, but does not contain any of the known ESCRT-binding motifs previously mentioned [88]. Therefore, it is difficult to predict the ESCRT-binding activity of a protein based on sequence information alone.

Previous studies have noted the knockdown of ESCRT components like the Vps2 homolog, CHMP2, and the SNF7 homolog, SHRB, in JW18 *Drosophila melanogaster* embryonic cell lines caused an increase in *Wolbachia* populations in vivo [89], indicating ESCRT activity likely plays an important role in *Wolbachia* intracellular persistence. Additionally, activation of autophagic pathways via exogenous addition of rapamycin to either *Wolbachia*-infected *Drosophila* cell lines or *Brugia malayi* was shown to reduce intracellular *Wolbachia* populations, presumably through lysosomal degradation of the bacterium [90]. As ESCRT activity is required for macro and microautophagy [91–93], it is possible that Wbm0152 (and orthologs) dampens host ESCRT activity to prevent autophagic degradation of the bacterium. Furthermore, RNA-Seq data obtained from 6-week *Brugia malayi* microfilaria isolated from tetracycline-treated animals to eliminate *Wolbachia* from the nematode, found that the transcription of *Brugia* genes producing proteins involved in multivesicular body formation were upregulated in the absence of *Wolbachia* [94], indicating that the presence of *Wolbachia* does appear to alter *Brugia*/filarial nematode endolysosomal membrane dynamics.

Interestingly, Wbm0152 belongs to a conserved family of peptidoglycan-associated lipoproteins (Pal) found extensively throughout gram negative bacteria, where it plays a role in bacterial outer membrane stability and in cell division [95, 96]. In Gram-negative bacteria, Pal proteins interact with the peptidoglycan cell wall, outer membrane proteins, and the periplasmic TolB protein – which is part of the larger inner membrane Tol complex (consisting of TolQ, TolR, and TolA) – and concentrates at the site of cell division to modulate peptidoglycan processing during cell division via the recruitment of cell separation amidases [97, 98]. While it is difficult to hypothesize how a periplasmic-facing outer membrane lipoprotein could interact with ‘external’ host cell proteins in the context of *Wolbachia* physiology, it is important to note that the *Wolbachia* genome does not appear to contain any known Tol protein homologs – including TolB [99] – suggesting the role of this Pal-like protein in *Wolbachia* may be quite different than typically observed in most Gram-negative bacteria. Furthermore, Pal-family proteins have been shown in some bacteria to have a ‘dual confirmation’ in which the population of Pal can have an ‘inward’ facing (periplasmic) C-terminus, or an ‘outward’ facing, exposed C-terminus on the outer membrane surface. Despite this well documented phenotype, the physiological role of this ‘outward’ facing Pal population remains unknown [100]. In support of the hypothesis that Wbm0152 is outward facing in *Wolbachia*, Wbm0152 has been previously localized to both the surface of the *Wolbachia* bacterium – as well as on the membrane of the *Wolbachia*-containing vacuole – via immunofluorescence and immuno-EM in *Brugia* tissues [101]. The fact that Wbm0152 is present on the surface of the *Wolbachia*-containing vacuole suggests that this protein may be positioned to interact directly with *Brugia* cytoplasmic proteins, like ESCRT.

Why might *Wolbachia* modulate ESCRT activity in *Brugia*? As *Wolbachia* lives inside of a membrane-bound compartment within nematode cells, we could hypothesize *Wolbachia* may be modulating *Brugia* ESCRT function to initiate ILV formation <u>into</u> the *Wolbachia*-containing compartment, perhaps to deliver nutrients from the cytosol of the host cell. Furthermore, this activity could also be utilized either for the creation of the *Wolbachia* containing vacuole from post-Golgi membranes, or to prevent entry into lysosomal degradation pathways that require ESCRT activity, such as autophagy. Lastly, it is known that *Wolbachia* is required to both maintain a quiescent pool of germline stem cells in the female *Brugia* germline and stimulates mitotic cell division during embryonic development [102]. As ESCRT-III activity was also found to be critical for abscission during cytokinesis and nuclear envelope repair during cell division [30, 103, 104], it is even possible that *Wolbachia* directly controls the cell cycle of the *Brugia* germline cells through the manipulation of ESCRT-III activity. Unfortunately, direct experiments to test these hypotheses are difficult without greatly expanding the availability of research reagents for both *Brugia* and *Wolbachia*. Nevertheless, this study marks an important milestone in the analysis of the molecular relationship between *Wolbachia* and the filarial nematode host, opening doors for new drug development for the eradication of *Wolbachia* – and subsequently – filarial nematodes.

## METHODS

### Yeast strain and plasmid constructions

All microscopy-based experiments were performed with derivatives of the yeast strain SEY6210 (*MATa ura3-52 leu2-3, 112 his3-Δ100 trp1-Δ901 lys2-801 suc2-Δ9*). For the growth experiments performed in Figs 2 and 7, yeast strains were derivatives of BY4742 (MATα his3Δ1 leu2Δ0 lys2Δ0 ura3Δ0). Any expression studies utilizing β-estradiol for the induction of *GAL1* promoters required transforming relevant yeast strains with linearized pAGL (a gift from Dr. Daniel Gottschling, University of Washington), thus introducing the gene encoding for the Gal4-estrogen receptor-VP16 (GEV) chimeric protein into the *leu2* locus [105].

To create a high-copy, constitutive yeast expression vector for *w*Bm0152, we replaced the *GAL1* promoter contained in pYES2/NT A (ThermoFisher Scientific) with the strong, constitutive *TDH3* promoter. The *TDH3* promoter was amplified from the p413-GPD plasmid [106] using the primer pair YTDH F and YTDHR **(Table S1**), each containing 30-bp of homology upstream and downstream of the *GAL1* promoter in pYES2/NT A. The resultant amplicon was co-transformed into BY4742 with pYES2/NT A that had previously been linearized with PvuII, using standard lithium acetate techniques [107]. Gap-repaired plasmids were selected on CSM medium lacking uracil. To generate the pYES*_TDH3_*-*w*Bm0152 plasmid, the pYES2-wBm0152 plasmid [19] was digested with PmeI and HindIII, the *w*Bm0152-containing insert was gel purified and co-transformed with pYES-*TDH3* previously linearized with BamHI, as above.

To create a high-copy, inducible yeast expression vector for *w*Bm0152 that does not rely on activation of the *GAL1* promoter, we replaced the *GAL1* promoter from pYES2/NT A with the copper-inducible *CUP1* promoter [108]. To avoid repetitive DNA sequences contained in the duplicated *S. cerevisiae CUP1-1 CUP1-2* locus, we first amplified the upstream sequence of *CUP1-1* with the primer pair CUP F and CUP R. The resultant amplicon was used as template with the primer pair YCUP F and YCUP R; the amplicon from this second reaction was then co-transformed into yeast with pYES2/NT A previously linearized with PvuII. To create the *w*Bm0152 expression construct in this background, pYES_CUP1_ was linearized with BamHI and co-transformed with the PmeI-HindIII *w*Bm0152-containing insert from above.

The copper-inducible pYES_CUP1_-*w*Bm0152 split Venus yeast expression plasmids used for the biomolecular fluorescence complementation assays were created by amplifying either the VN or VC domain from the relevant plasmid template using the indicated primer pairs containing 30-bp homology to the C-terminus of wBm0152 and the pYES plasmid backbone (VN: pFA6a-VN-HIS3 with 0152VCN F and 0152VN R; VC: pFA6a-VC-TRP1 with 0152VCN F and 0152VC R) [62]. The resultant amplicons were separately co-transformed into yeast with PmeI-linearized pYES*_CUP1_*-*w*Bm0152 to create the final constructs via gap repair. To genera_te t_he pYES_CUP1_-*VPS2-*VN split Venus yeast expression plasmid, the VPS2-VN fusion was amplified off the genome of the SEY6210 *VPS2-*VN yeast strain, using the primer pair VPS2VN F and VPS2VN R. The resultant amplicon was co-transformed into yeast with BamHI-linearized pYES*_CUP1_*-*w*Bm0152 to create the final construct via gap repair.

The *Brugia malayi VPS2* homolog was identified via blastp [109] using the yeast Did4p/Vps2p sequence and restricting the search to *Brugia malayi*. The closest match was *Bm6583b* (GenBank accession: VIO93588) and the transcript cDNA encoding for this ORF was retrieved from the WormBase web site, http://www.wormbase.org, <u>release WS296,</u> <u>date Apr 30 2025</u> [110]. This sequence was codon-optimized for *Saccharomyces cerevisiae* expression using the Codon Optimization Tool (IDT™ Technologies). When generating the *Bm6583b* gBlock®(IDT™ Technologies), we also appended the 300 base pairs immediately upstream of the yeast *VPS2* ORF, containing the endogenous promoter region. Prior to the *Bm6583b* stop codon, we added sequences encoding for the c-Myc epitope for detection via immunoblot, a BamHI restriction site immediately after the stop codon, and the 300 bases immediately downstream of the yeast *VPS2* ORF, containing the endogenous terminator region. Finally, we appended 30-bp sequences to the 5’ and 3’ ends of this sequence, providing homology flanking the MCS of the yeast plasmid pRS414 for cloning via gap repair in yeast (**Table S1**). This gBlock® was co-transformed into yeast with pRS414 previously linearized with BamHI, creating pRS414-*Bm6583*-myc. To generate the *Bm6583* expression plasmid used for the split Venus bimolecular fluorescence complementation study, the Venus VN domain was amplified from pFA6a-VN-HIS3 with the primer pair *BM6583* VN F and *BM6583* VN R (**Table S1**) and co-transformed into yeast with pRS414-*Bm6583-*myc plasmid previously digested with BamHI.

All generated plasmid constructs were confirmed through whole plasmid sequencing by Plasmidsaurus using Oxford Nanopore Technology with custom analysis and annotation.

### RapIDeg protein turnover assay

To visualize the ESCRT-dependent turnover of Fth1p in response to rapamycin, we modified the RapIDeg yeast strain Fth1-GFP-FKBP 3x Ub (SEY6210.1 *tor1-1 fpr1Δ::NATMX6* pRS305-pGPD-FRB-3xUb::*LEU2* FTH1-GFP-2xFKBP::*HIS3*, a gift from Dr. Ming Li, University of Michigan) [48] to express *GAL* promoters with β-estradiol. First, we amplified the *HPHMX6* gene from pAG32 [111] using primer pair pr29 and pr32 (**Table S1**); this amplicon contains homology to the *NATMX6* cassette inserted into the *FPR1* gene. The RapIDeg strain was transformed with this amplicon and hygromycin-resistant colonies were selected. This strain was then screened for nourseothricin sensitivity, confirming the replacement of the *NATMX6* cassette with *HPHMX6*. The resultant strain was then transformed with the linear pAGL, as above, creating the β-estradiol-responsive RapIDeg strain.

RapIDeg strains harboring either the galactose-inducible pYES-*w*Bm0152 plasmid or empty vector control were grown to saturation in selective medium at 30° C for 16h. Strains were subcultured into 10 mL fresh selective medium and grown to mid-log for 4h at 30 °C. To induce the expression of *w*Bm0152, 1 µM β-estradiol was added to each culture and incubated at 30° C with shaking for another 2h. After 2h, each culture was evenly split and either rapamycin or the DMSO vehicle control was added to 1 µg mL^−1^ and cultures were outgrown for an additional 2h at 30°. 1 mL aliquots were removed from each condition, yeast cells were harvested by centrifugation (7000 x *g,* 1 min), and visualized via fluorescence microscopy. From the remaining culture, 3.0 OD_600_ units were harvested via centrifugation and total cellular proteins were extracted from cell pellets [112] for immunoblotting experiments.

### Microscopy

Cells to be visualized were grown overnight at 30° C in selective media, subcultured in fresh media with or without induction agent(s), and grown for an additional 5 hours. Cells were harvested via centrifugation, washed with sterile water, and suspended in 50 µL water. Cell suspensions were mounted to slides pre-treated with a 1:1 mixture of polylysine (10% w/v):concanavalin A (2 mg mL^−1^) solution. Cells were visualized using a Nikon Ti-U fluorescence microscope, and images were processed using the Fiji software package (ImageJ2, v2.14.0/1.54f) [113, 114]. Colocalization analyses were performed using the Coloc2 plugin contained within the Fiji package.

### Thin section electron microscopy

*Saccharomyces cerevisiae* strain SEY6210 *GAL^+^* harboring either a vector control or the pYES-*w*Bm0152 expression plasmid were grown in selective media (CSM lacking uracil) for 18h at 30° C with shaking. Cells were harvested via centrifugation, washed with sterile water, then diluted 1:10 into fresh CSM-ura containing 2% galactose to induce *w*Bm0152 expression. Cells were harvested at log phase, collected on 0.45µ filter discs (Millipore) via vacuum, loaded into 0.25 mm aluminum hats, and high-pressure frozen in the Wolwend high-pressure freezing machine (HPF) as previously described [56]. Freeze substitution was carried out with media containing 0.1% uranyl acetate and 0.25% gluteraldehyde in anhydrous acetone [115] in a Leica AFS (Automated Freeze Substitution, Vienna, Austria). Samples were then embedded in Lowicryl HM20, and UV-polymerized at - 60C over 2 weeks (Polysciences, Warrington, PA, A. Staehelin, personal communication, Wemmer et al JCB 2011). A Leica Ultra-Microtome was used to cut 80 nm serial thin sections and 200 nm serial semi-thick sections from polymerized blocks and sections were collected onto 1% formvar films adhered to rhodium-plated copper grids (Leica Biosystems, Nussloch, Germany, and Electron Microscopy Sciences). Thin sections were imaged using a FEI Tecnai T12 “Spirit” electron microscope (120 kV, AMT 2 x 2 k CCD).

### Semi-thick section electron tomography

Semi-thick sections of polymerized blocks (∼200 nm) were applied to grids labeled on both sides with fiduciary 15 nm colloidal gold (British Biocell International). Dual-axis tilt series were collected of the samples from +/− 60 degrees with 1-degree increments at 300kV using SerialEM [116] on a Tecnai 30 FEG electron microscope (FEI-Company, Eindhoven, the Netherlands). Tilt series were regularly imaged at 23,000X using SerialEM [117], with 2X binning when recording on a 4×4K CCD camera (Gatan, Inc., Abingdon, UK) creating a 2×2K image with a pixel size of 1.02 nm. Tomograms were constructed and modeled with the IMOD software package (3dmod 4.0.11, [118]). All membrane structures were identified with areas of interest modeled (ER-LD compartments, small MVBs, tubular MVBs). IMODINFO provided surface area and volume data of contour models, diameters and distances were measured from the outer membrane leaflets at optimal X Y orientations in the tomograms at 50 nm intervals using 3DMOD.

## Acknowledgments

The authors would like to thank Dr. Ming Li (University of Michigan) for providing the yeast RapIDeg strain used in these studies. V.J.S. is supported by a grant from the National Institute of Allergy and Infectious Diseases (R21-AI171573). G.O. is supported by a grant from the National Institute of General Medical Sciences (R32 GM149202).

## Supplemental Information

**Figure S1.**
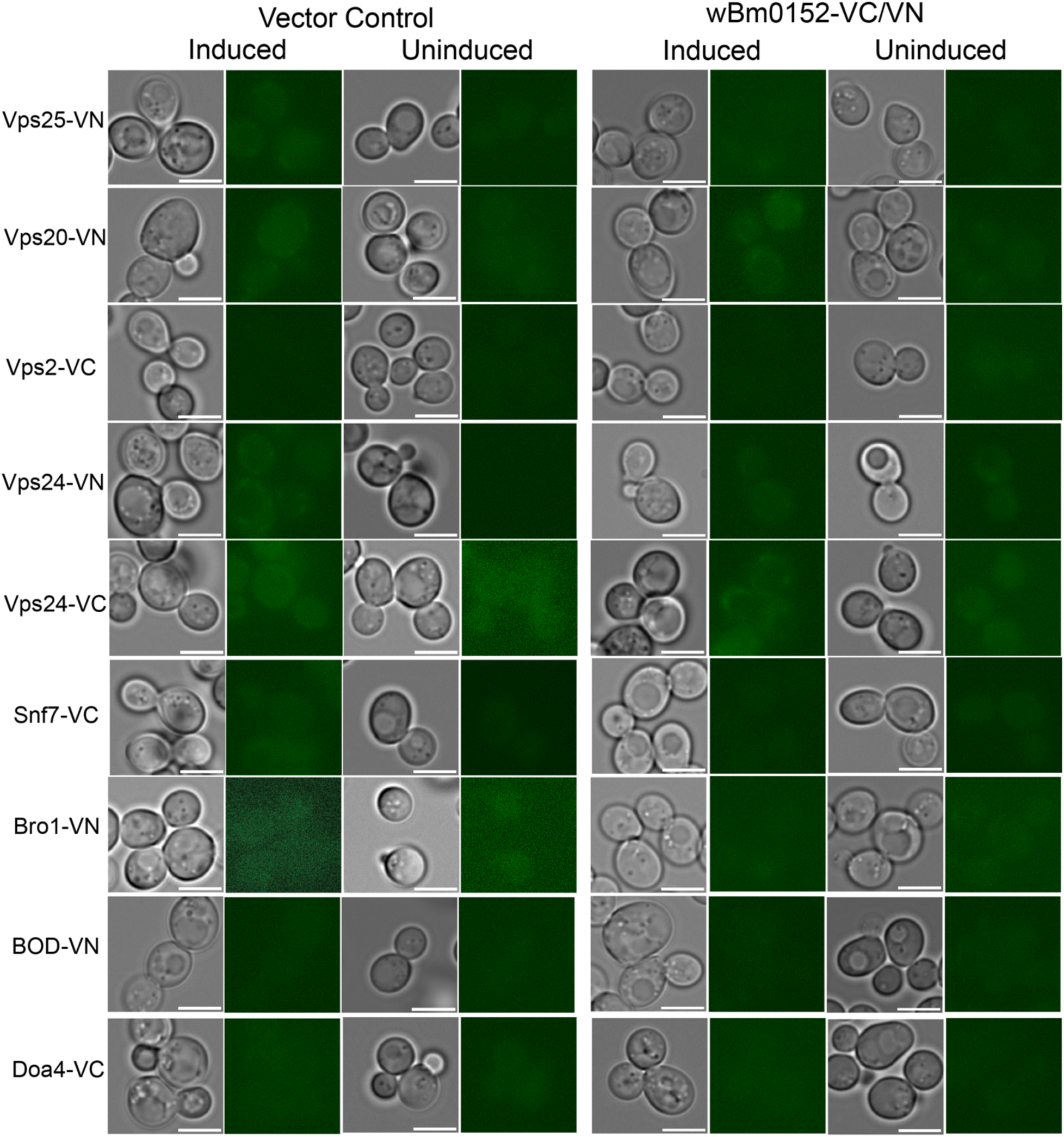
wBm0152 does not bind other ESCRT subunits in vivo. SEY6210 strains harboring either the N-terminus (VN) or C-terminus (VC) of a Venus-YFP molecule on the C-terminus of the indicated ESCRT subunit were transformed with the corresponding copper-inducible pYES*_CUP1_*-*w*Bm0152-VN or pYES*_CUP1_-w*Bm0152-VC plasmid. Strains were grown in CSM media lacking uracil for 18h at 30°C with shaking, diluted 1:10 into fresh selective media lacking or supplemented with 0.5 mM CuSO_4_, and outgrown for 6 hours before imaging. Bar = 5 µ; images are representative of three separate experiments.

**Table S1.**
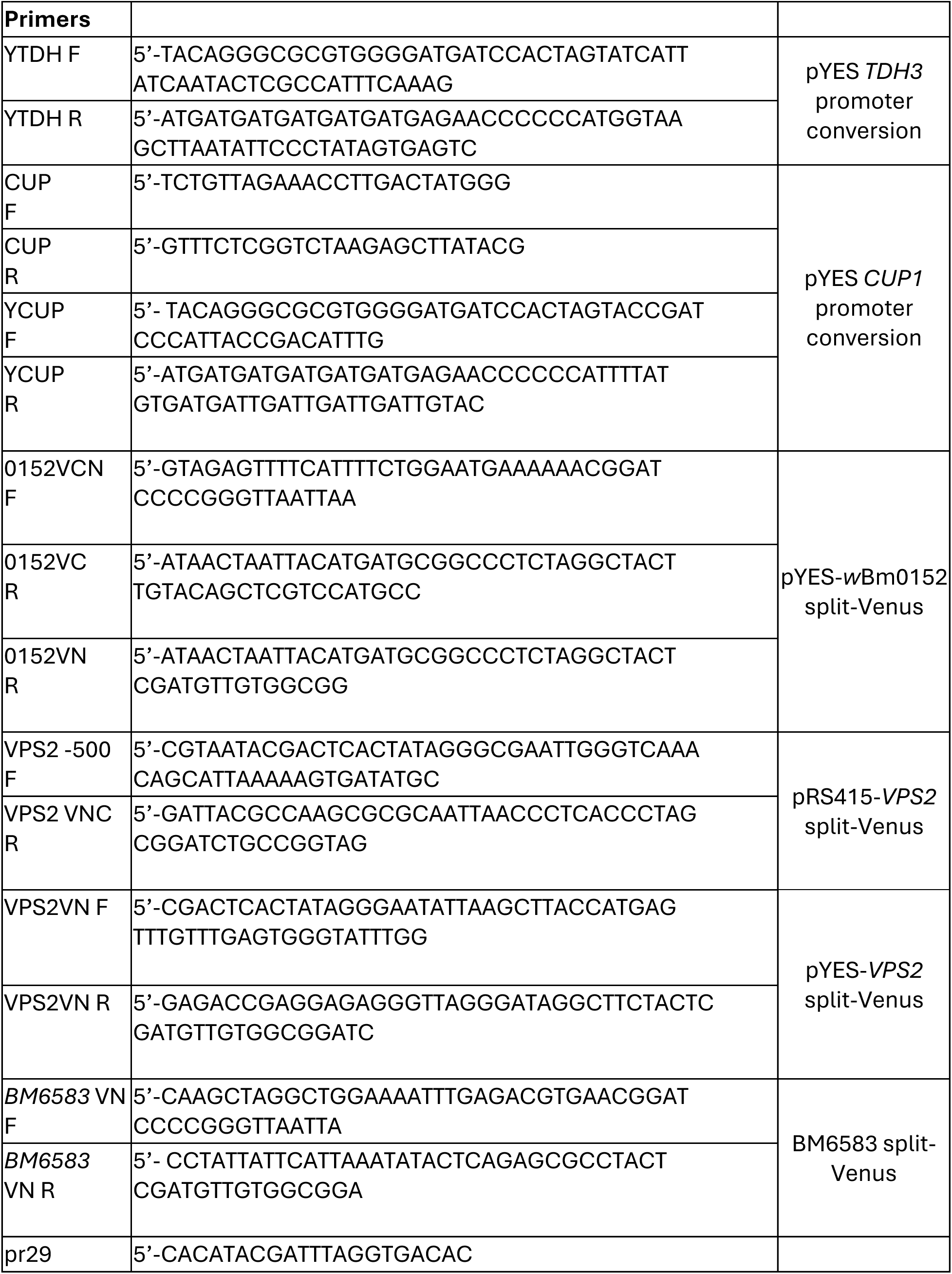
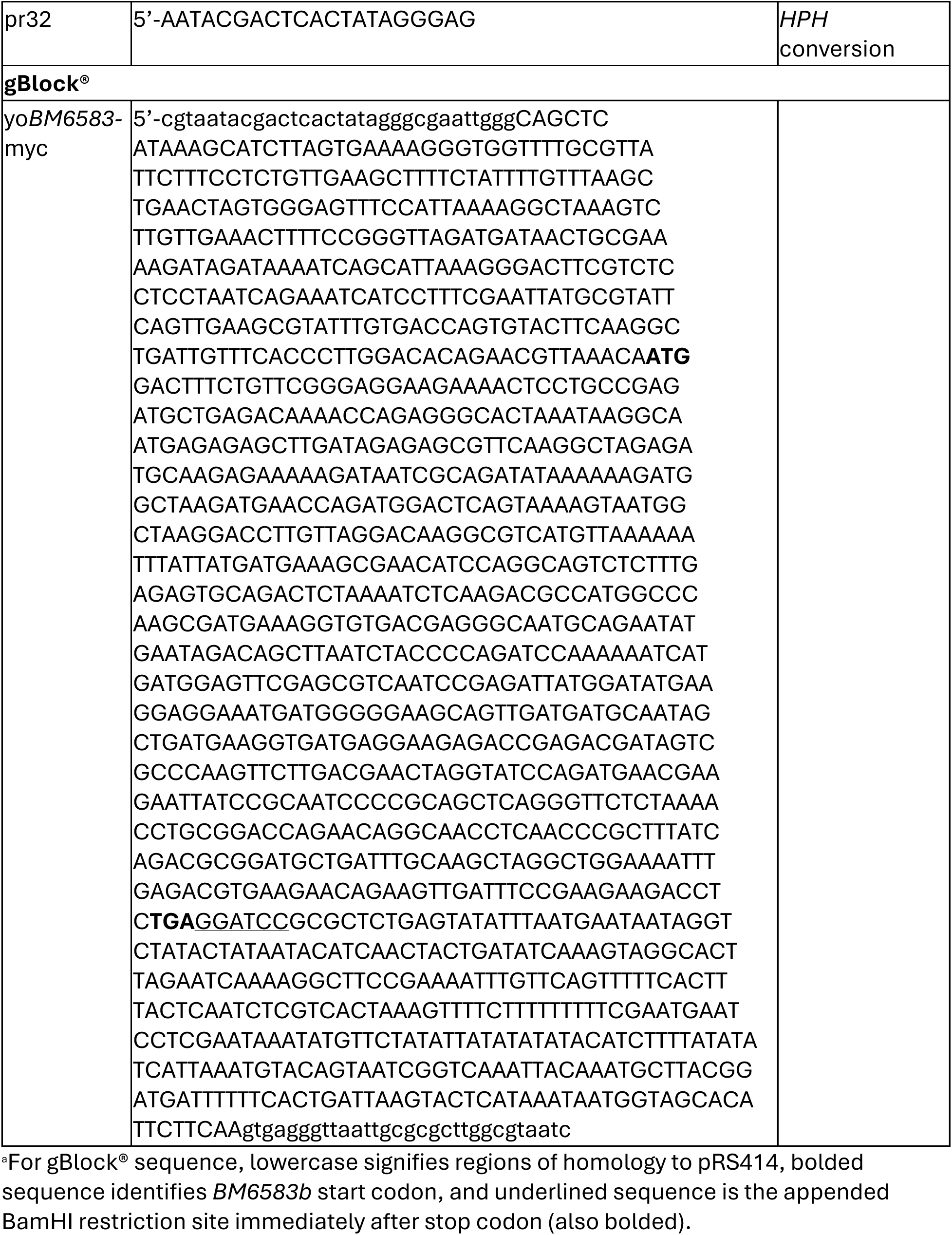
Primer and nucleotide sequences used in this study^a^.

## References

1. Aziz MA, Diallo S, Diop IM, Lariviere M, Porta M. Efficacy and tolerance of ivermectin in human onchocerciasis. Lancet. 1982;2(8291):171–3. doi: 10.1016/s0140-6736(82)91026-1. PubMed PMID: 6123884.

2. Mak JW, Navaratnam V, Grewel JS, Mansor SM, Ambu S. Treatment of subperiodic *Brugia malayi* infection with a single dose of ivermectin. Am J Trop Med Hyg. 1993;48(4):591–6. doi: 10.4269/ajtmh.1993.48.591. PubMed PMID: 8480868.

3. Shenoy RK, John A, Babu BS, Suma TK, Kumaraswami V. Two-year follow-up of the microfilaraemia of asymptomatic brugian filariasis, after treatment with two, annual, single doses of ivermectin, diethylcarbamazine and albendazole, in various combinations. Ann Trop Med Parasitol. 2000;94(6):607–14. doi: 10.1080/00034983.2000.11813583. PubMed PMID: 11064762.

4. Campbell WC. Ivermectin as an antiparasitic agent for use in humans. Annu Rev Microbiol. 1991;45:445–74. doi: 10.1146/annurev.mi.45.100191.002305. PubMed PMID: 1741621.

5. Dreyer G, Addiss D, Santos A, Figueredo-Silva J, Noroes J. Direct assessment in vivo of the efficacy of combined single-dose ivermectin and diethylcarbamazine against adult *Wuchereria bancrofti*. Trans R Soc Trop Med Hyg. 1998;92(2):219–22. doi: 10.1016/s0035-9203(98)90754-4. PubMed PMID: 9764338.

6. Kashyap SS, Verma S, McHugh M, Wolday M, Williams PD, Robertson AP, et al. Anthelmintic resistance and homeostatic plasticity (Brugia malayi). Sci Rep. 2021;11(1):14499. Epub 20210714. doi: 10.1038/s41598-021-93911-4. PubMed PMID: 34262123; PubMed Central PMCID: PMC8280109.

7. Osei-Atweneboana MY, Awadzi K, Attah SK, Boakye DA, Gyapong JO, Prichard RK. Phenotypic evidence of emerging ivermectin resistance in *Onchocerca volvulus*. PLoS Negl Trop Dis. 2011;5(3):e998. doi: 10.1371/journal.pntd.0000998. PubMed PMID: 21468315; PubMed Central PMCID: PMC3066159.

8. Osei-Atweneboana MY, Boakye DA, Awadzi K, Gyapong JO, Prichard RK. Genotypic analysis of beta-tubulin in *Onchocerca volvulus* from communities and individuals showing poor parasitological response to ivermectin treatment. Int J Parasitol Drugs Drug Resist. 2012;2:20–8. doi: 10.1016/j.ijpddr.2012.01.005. PubMed PMID: 24533268; PubMed Central PMCID: PMC3862422.

9. Sironi M, Bandi C, Sacchi L, Di Sacco B, Damiani G, Genchi C. Molecular evidence for a close relative of the arthropod endosymbiont *Wolbachia* in a filarial worm. Mol Biochem Parasitol. 1995;74(2):223–7. doi: 10.1016/0166-6851(95)02494-8. PubMed PMID: 8719164.

10. Taylor MJ, Bandi C, Hoerauf A. *Wolbachia* bacterial endosymbionts of filarial nematodes. Adv Parasitol. 2005;60:245–84. doi: 10.1016/S0065-308X(05)60004-8. PubMed PMID: 16230105.

11. Debrah AY, Mand S, Marfo-Debrekyei Y, Batsa L, Albers A, Specht S, et al. Macrofilaricidal Activity in *Wuchereria bancrofti* after 2 Weeks Treatment with a Combination of Rifampicin plus Doxycycline. J Parasitol Res. 2011;2011:201617. doi: 10.1155/2011/201617. PubMed PMID: 21687646; PubMed Central PMCID: PMC3112504.

12. Debrah AY, Mand S, Marfo-Debrekyei Y, Batsa L, Pfarr K, Buttner M, et al. Macrofilaricidal effect of 4 weeks of treatment with doxycycline on *Wuchereria bancrofti*. Trop Med Int Health. 2007;12(12):1433–41. doi: 10.1111/j.1365-3156.2007.01949.x. PubMed PMID: 18076549.

13. Landmann F, Voronin D, Sullivan W, Taylor MJ. Anti-filarial activity of antibiotic therapy is due to extensive apoptosis after *Wolbachia* depletion from filarial nematodes. PLoS Pathog. 2011;7(11):e1002351. doi: 10.1371/journal.ppat.1002351. PubMed PMID: 22072969; PubMed Central PMCID: PMC3207916.

14. Taylor MJ, Makunde WH, McGarry HF, Turner JD, Mand S, Hoerauf A. Macrofilaricidal activity after doxycycline treatment of *Wuchereria bancrofti*: a double-blind, randomised placebo-controlled trial. Lancet. 2005;365(9477):2116-21. doi: 10.1016/S0140-6736(05)66591-9. PubMed PMID: 15964448.

15. Duclos S, Desjardins M. Subversion of a young phagosome: the survival strategies of intracellular pathogens. Cell Microbiol. 2000;2(5):365–77. doi: 10.1046/j.1462-5822.2000.00066.x. PubMed PMID: 11207592.

16. Huang B, Hubber A, McDonough JA, Roy CR, Scidmore MA, Carlyon JA. The *Anaplasma phagocytophilum*-occupied vacuole selectively recruits Rab-GTPases that are predominantly associated with recycling endosomes. Cell Microbiol. 2010;12(9):1292–307. Epub 20100325. doi: 10.1111/j.1462-5822.2010.01468.x. PubMed PMID: 20345488; PubMed Central PMCID: PMC2923681.

17. Cho KO, Kim GW, Lee OK. *Wolbachia* bacteria reside in host Golgi-related vesicles whose position is regulated by polarity proteins. PLoS One. 2011;6(7):e22703. doi: 10.1371/journal.pone.0022703. PubMed PMID: 21829485; PubMed Central PMCID: PMC3145749.

18. Fattouh N, Cazevieille C, Landmann F. *Wolbachia* endosymbionts subvert the endoplasmic reticulum to acquire host membranes without triggering ER stress. PLoS Negl Trop Dis. 2019;13(3):e0007218. Epub 2019/03/21. doi: 10.1371/journal.pntd.0007218. PubMed PMID: 30893296; PubMed Central PMCID: PMC6426186.

19. Carpinone EM, Li Z, Mills MK, Foltz C, Brannon ER, Carlow CKS, et al. Identification of putative effectors of the Type IV secretion system from the *Wolbachia* endosymbiont of *Brugia malayi*. PLoS One. 2018;13(9):e0204736. Epub 2018/09/28. doi: 10.1371/journal.pone.0204736. PubMed PMID: 30261054; PubMed Central PMCID: PMC6160203.

20. Li Z, Carlow CK. Characterization of transcription factors that regulate the type IV secretion system and riboflavin biosynthesis in *Wolbachia* of *Brugia malayi*. PLoS One. 2012;7(12):e51597. doi: 10.1371/journal.pone.0051597. PubMed PMID: 23251587; PubMed Central PMCID: PMC3518464.

21. Rances E, Voronin D, Tran-Van V, Mavingui P. Genetic and functional characterization of the type IV secretion system in *Wolbachia*. J Bacteriol. 2008;190(14):5020–30. doi: 10.1128/JB.00377-08. PubMed PMID: 18502862; PubMed Central PMCID: PMC2447017.

22. Whitaker N, Berry TM, Rosenthal N, Gordon JE, Gonzalez-Rivera C, Sheehan KB, et al. Chimeric Coupling Proteins Mediate Transfer of Heterologous Type IV Effectors through the *Escherichia coli* pKM101-Encoded Conjugation Machine. J Bacteriol. 2016;198(19):2701–18. Epub 2016/07/20. doi: 10.1128/JB.00378-16. PubMed PMID: 27432829; PubMed Central PMCID: PMC5019051.

23. Mills MK, McCabe LG, Rodrigue EM, Lechtreck KF, Starai VJ. Wbm0076, a candidate effector protein of the *Wolbachia* endosymbiont of *Brugia malayi*, disrupts eukaryotic actin dynamics. PLoS Pathog. 2023;19(2):e1010777. Epub 20230217. doi: 10.1371/journal.ppat.1010777. PubMed PMID: 36800397; PubMed Central PMCID: PMC9980815.

24. Babst M. MVB vesicle formation: ESCRT-dependent, ESCRT-independent and everything in between. Curr Opin Cell Biol. 2011;23(4):452–7. doi: 10.1016/j.ceb.2011.04.008. PubMed PMID: 21570275; PubMed Central PMCID: PMC3148405.

25. Gruenberg J, Stenmark H. The biogenesis of multivesicular endosomes. Nat Rev Mol Cell Biol. 2004;5(4):317–23. doi: 10.1038/nrm1360. PubMed PMID: 15071556.

26. Katzmann DJ, Odorizzi G, Emr SD. Receptor downregulation and multivesicular-body sorting. Nat Rev Mol Cell Biol. 2002;3(12):893–905. doi: 10.1038/nrm973. PubMed PMID: 12461556.

27. Oku M, Maeda Y, Kagohashi Y, Kondo T, Yamada M, Fujimoto T, et al. Evidence for ESCRT- and clathrin-dependent microautophagy. J Cell Biol. 2017;216(10):3263–74. Epub 20170824. doi: 10.1083/jcb.201611029. PubMed PMID: 28838958; PubMed Central PMCID: PMC5626533.

28. Raiborg C, Stenmark H. The ESCRT machinery in endosomal sorting of ubiquitylated membrane proteins. Nature. 2009;458(7237):445–52. doi: 10.1038/nature07961. PubMed PMID: 19325624.

29. Jimenez AJ, Maiuri P, Lafaurie-Janvore J, Divoux S, Piel M, Perez F. ESCRT machinery is required for plasma membrane repair. Science. 2014;343(6174):1247136. Epub 20140130. doi: 10.1126/science.1247136. PubMed PMID: 24482116.

30. Olmos Y, Hodgson L, Mantell J, Verkade P, Carlton JG. ESCRT-III controls nuclear envelope reformation. Nature. 2015;522(7555):236–9. Epub 20150603. doi: 10.1038/nature14503. PubMed PMID: 26040713; PubMed Central PMCID: PMC4471131.

31. Raab M, Gentili M, de Belly H, Thiam HR, Vargas P, Jimenez AJ, et al. ESCRT III repairs nuclear envelope ruptures during cell migration to limit DNA damage and cell death. Science. 2016;352(6283):359–62. Epub 20160324. doi: 10.1126/science.aad7611. PubMed PMID: 27013426.

32. Baumgartel V, Ivanchenko S, Dupont A, Sergeev M, Wiseman PW, Krausslich HG, et al. Live-cell visualization of dynamics of HIV budding site interactions with an ESCRT component. Nat Cell Biol. 2011;13(4):469–74. Epub 20110310. doi: 10.1038/ncb2215. PubMed PMID: 21394086.

33. Morita E, Sandrin V, McCullough J, Katsuyama A, Baci Hamilton I, Sundquist WI. ESCRT-III protein requirements for HIV-1 budding. Cell Host Microbe. 2011;9(3):235–42. doi: 10.1016/j.chom.2011.02.004. PubMed PMID: 21396898; PubMed Central PMCID: PMC3070458.

34. Pincetic A, Medina G, Carter C, Leis J. Avian sarcoma virus and human immunodeficiency virus, type 1 use different subsets of ESCRT proteins to facilitate the budding process. J Biol Chem. 2008;283(44):29822–30. Epub 20080822. doi: 10.1074/jbc.M804157200. PubMed PMID: 18723511; PubMed Central PMCID: PMC2573067.

35. Banjade S, Tang S, Shah YH, Emr SD. Electrostatic lateral interactions drive ESCRT-III heteropolymer assembly. Elife. 2019;8. Epub 20190627. doi: 10.7554/eLife.46207. PubMed PMID: 31246173; PubMed Central PMCID: PMC6663469.

36. Henne WM, Buchkovich NJ, Zhao Y, Emr SD. The endosomal sorting complex ESCRT-II mediates the assembly and architecture of ESCRT-III helices. Cell. 2012;151(2):356–71. doi: 10.1016/j.cell.2012.08.039. PubMed PMID: 23063125.

37. Hurley JH, Emr SD. The ESCRT complexes: structure and mechanism of a membrane-trafficking network. Annu Rev Biophys Biomol Struct. 2006;35:277–98. doi: 10.1146/annurev.biophys.35.040405.102126. PubMed PMID: 16689637; PubMed Central PMCID: PMC1648078.

38. Jouvenet N. Dynamics of ESCRT proteins. Cell Mol Life Sci. 2012;69(24):4121–33. Epub 20120606. doi: 10.1007/s00018-012-1035-0. PubMed PMID: 22669260; PubMed Central PMCID: PMC11114710.

39. Tang S, Henne WM, Borbat PP, Buchkovich NJ, Freed JH, Mao Y, et al. Structural basis for activation, assembly and membrane binding of ESCRT-III Snf7 filaments. Elife. 2015;4. Epub 20151215. doi: 10.7554/eLife.12548. PubMed PMID: 26670543; PubMed Central PMCID: PMC4720517.

40. Teis D, Saksena S, Emr SD. Ordered assembly of the ESCRT-III complex on endosomes is required to sequester cargo during MVB formation. Dev Cell. 2008;15(4):578–89. doi: 10.1016/j.devcel.2008.08.013. PubMed PMID: 18854142.

41. Babst M, Wendland B, Estepa EJ, Emr SD. The Vps4p AAA ATPase regulates membrane association of a Vps protein complex required for normal endosome function. EMBO J. 1998;17(11):2982–93. doi: 10.1093/emboj/17.11.2982. PubMed PMID: 9606181; PubMed Central PMCID: PMC1170638.

42. Buysse D, Pfitzner AK, West M, Roux A, Odorizzi G. The ubiquitin hydrolase Doa4 directly binds Snf7 to inhibit recruitment of ESCRT-III remodeling factors in *S. cerevisiae*. J Cell Sci. 2020;133(8). Epub 20200428. doi: 10.1242/jcs.241455. PubMed PMID: 32184262; PubMed Central PMCID: PMC7197871.

43. Richter CM, West M, Odorizzi G. Doa4 function in ILV budding is restricted through its interaction with the Vps20 subunit of ESCRT-III. J Cell Sci. 2013;126(Pt 8):1881–90. Epub 20130226. doi: 10.1242/jcs.122499. PubMed PMID: 23444383; PubMed Central PMCID: PMC3678411.

44. Raymond CK, Howald-Stevenson I, Vater CA, Stevens TH. Morphological classification of the yeast vacuolar protein sorting mutants: evidence for a prevacuolar compartment in class E vps mutants. Mol Biol Cell. 1992;3(12):1389–402. PubMed PMID: 1493335; PubMed Central PMCID: PMC275707.

45. Zhu L, Jorgensen JR, Li M, Chuang YS, Emr SD. ESCRTs function directly on the lysosome membrane to downregulate ubiquitinated lysosomal membrane proteins. Elife. 2017;6. Epub 20170629. doi: 10.7554/eLife.26403. PubMed PMID: 28661397; PubMed Central PMCID: PMC5507667.

46. Katzmann DJ, Sarkar, S., Chu, T., Audhya A., Emr, S.D. Multivesicular Body Sorting: Ubiquitin Ligase Rsp5 Is Required for the Modification and Sorting of Carboxypeptidase S. Molecular Biology of the Cell. 2004;15:468–80. doi: 10.1091/mbc.E03-.

47. Odorizzi G, Babst M, Emr SD. Fab1p PtdIns(3)P 5-kinase function essential for protein sorting in the multivesicular body. Cell. 1998;95(6):847–58. PubMed PMID: 9865702.

48. Yang X, Reist L, Chomchai DA, Chen L, Arines FM, Li M. ESCRT, not intralumenal fragments, sorts ubiquitinated vacuole membrane proteins for degradation. J Cell Biol. 2021;220(8). Epub 20210528. doi: 10.1083/jcb.202012104. PubMed PMID: 34047770; PubMed Central PMCID: PMC8167898.

49. Kopecka M, Gabriel M. The influence of congo red on the cell wall and (1 3)-beta-D-glucan microfibril biogenesis in *Saccharomyces cerevisiae*. Arch Microbiol. 1992;158(2):115–26. doi: 10.1007/BF00245214. PubMed PMID: 1417414.

50. Brune T, Kunze-Schumacher H, Kolling R. Interactions in the ESCRT-III network of the yeast *Saccharomyces cerevisiae*. Curr Genet. 2019;65(2):607–19. Epub 20181201. doi: 10.1007/s00294-018-0915-8. PubMed PMID: 30506264.

51. Luhtala N, Odorizzi G. Bro1 coordinates deubiquitination in the multivesicular body pathway by recruiting Doa4 to endosomes. J Cell Biol. 2004;166(5):717–29. Epub 20040823. doi: 10.1083/jcb.200403139. PubMed PMID: 15326198; PubMed Central PMCID: PMC2172414.

52. Wemmer M, Azmi I, West M, Davies B, Katzmann D, Odorizzi G. Bro1 binding to Snf7 regulates ESCRT-III membrane scission activity in yeast. J Cell Biol. 2011;192(2):295–306. doi: 10.1083/jcb.201007018. PubMed PMID: 21263029; PubMed Central PMCID: PMC3172170.

53. Buysse D, West M, Leih M, Odorizzi G. Bro1 binds the Vps20 subunit of ESCRT-III and promotes ESCRT-III regulation by Doa4. Traffic. 2022;23(2):109–19. Epub 20220113. doi: 10.1111/tra12828. PubMed PMID: 34908216; PubMed Central PMCID: PMC8792227.

54. Johnson N, West M, Odorizzi G. Regulation of yeast ESCRT-III membrane scission activity by the Doa4 ubiquitin hydrolase. Mol Biol Cell. 2017;28(5):661–72. Epub 20170105. doi: 10.1091/mbc.E16-11-0761. PubMed PMID: 28057764; PubMed Central PMCID: PMC5328624.

55. Nickerson DP, West M, Henry R, Odorizzi G. Regulators of Vps4 ATPase activity at endosomes differentially influence the size and rate of formation of intralumenal vesicles. Mol Biol Cell. 2010;21(6):1023–32. Epub 20100120. doi: 10.1091/mbc.e09-09-0776. PubMed PMID: 20089837; PubMed Central PMCID: PMC2836955.

56. Nickerson DP, West M, Odorizzi G. Did2 coordinates Vps4-mediated dissociation of ESCRT-III from endosomes. J Cell Biol. 2006;175(5):715–20. Epub 20061127. doi: 10.1083/jcb.200606113. PubMed PMID: 17130288; PubMed Central PMCID: PMC2064671.

57. Pfitzner AK, Zivkovic H, Bernat-Silvestre C, West M, Peltier T, Humbert F, et al. Vps60 initiates alternative ESCRT-III filaments. J Cell Biol. 2023;222(11). Epub 20230928. doi: 10.1083/jcb.202206028. PubMed PMID: 37768378; PubMed Central PMCID: PMC10538557.

58. Tseng CC, Dean S, Davies BA, Azmi IF, Pashkova N, Payne JA, et al. Bro1 stimulates Vps4 to promote intralumenal vesicle formation during multivesicular body biogenesis. J Cell Biol. 2021;220(8). Epub 20210623. doi: 10.1083/jcb.202102070. PubMed PMID: 34160559; PubMed Central PMCID: PMC8240856.

59. Babst M, Katzmann DJ, Snyder WB, Wendland B, Emr SD. Endosome-associated complex, ESCRT-II, recruits transport machinery for protein sorting at the multivesicular body. Dev Cell. 2002;3(2):283–9. doi: 10.1016/s1534-5807(02)00219-8. PubMed PMID: 12194858.

60. Katzmann DJ, Stefan CJ, Babst M, Emr SD. Vps27 recruits ESCRT machinery to endosomes during MVB sorting. J Cell Biol. 2003;162(3):413–23. doi: 10.1083/jcb.200302136. PubMed PMID: 12900393; PubMed Central PMCID: PMC2172707.

61. Ghosh I, Hamilton AD, Regan L. Antiparallel Leucine Zipper-Directed Protein Reassembly: Application to the Green Fluorescent Protein. Journal of the American Chemical Society. 2000;122(23):5658–9. doi: 10.1021/ja994421w.

62. Shyu YJ, Liu H, Deng X, Hu CD. Identification of new fluorescent protein fragments for bimolecular fluorescence complementation analysis under physiological conditions. Biotechniques. 2006;40(1):61–6. doi: 10.2144/000112036. PubMed PMID: 16454041.

63. Kerppola TK. Design and implementation of bimolecular fluorescence complementation (BiFC) assays for the visualization of protein interactions in living cells. Nat Protoc. 2006;1(3):1278–86. doi: 10.1038/nprot.2006.201. PubMed PMID: 17406412; PubMed Central PMCID: PMC2518326.

64. Strack B, Calistri A, Craig S, Popova E, Gottlinger HG. AIP1/ALIX is a binding partner for HIV-1 p6 and EIAV p9 functioning in virus budding. Cell. 2003;114(6):689–99. doi: 10.1016/s0092-8674(03)00653-6. PubMed PMID: 14505569.

65. Zamborlini A, Usami Y, Radoshitzky SR, Popova E, Palu G, Gottlinger H. Release of autoinhibition converts ESCRT-III components into potent inhibitors of HIV-1 budding. Proc Natl Acad Sci U S A. 2006;103(50):19140–5. Epub 20061204. doi: 10.1073/pnas.0603788103. PubMed PMID: 17146056; PubMed Central PMCID: PMC1748189.

66. Gill DJ, Teo H, Sun J, Perisic O, Veprintsev DB, Emr SD, et al. Structural insight into the ESCRT-I/-II link and its role in MVB trafficking. EMBO J. 2007;26(2):600–12. Epub 20070111. doi: 10.1038/sj.emboj.7601501. PubMed PMID: 17215868; PubMed Central PMCID: PMC1783442.

67. Teo H, Gill DJ, Sun J, Perisic O, Veprintsev DB, Vallis Y, et al. ESCRT-I core and ESCRT-II GLUE domain structures reveal role for GLUE in linking to ESCRT-I and membranes. Cell. 2006;125(1):99–111. doi: 10.1016/j.cell.2006.01.047. PubMed PMID: 16615893.

68. Ghedin E, Wang S, Spiro D, Caler E, Zhao Q, Crabtree J, et al. Draft genome of the filarial nematode parasite *Brugia malayi*. Science. 2007;317(5845):1756–60. Epub 2007/09/22. doi: 10.1126/science.1145406. PubMed PMID: 17885136; PubMed Central PMCID: PMC2613796.

69. Nevalainen LB, Layton EM, Newton ILG. *Wolbachia* Promotes Its Own Uptake by Host Cells. Infect Immun. 2023;91(2):e0055722. Epub 20230117. doi: 10.1128/iai.00557-22. PubMed PMID: 36648231; PubMed Central PMCID: PMC9933726.

70. Newton ILG, Savytskyy O., Sheehan, K.B. *Wolbachia* Utilize Host Actin for Efficient Maternal Transmission in *Drosophila melanogaster*. PLoS Pathog. 2015;11(4):e1004798. doi: 10.1371/journal.ppat.1004798.

71. Rice DW, Sheehan, K.B., Newton, I.L.G. Large-Scale Identification of *Wolbachia pipientis* Effectors. Genome Biol Evol 2017;9(7):1925–37.

72. Lesser CF, Miller SI. Expression of microbial virulence proteins in *Saccharomyces cerevisiae* models mammalian infection. EMBO J. 2001;20(8):1840–9. doi: 10.1093/emboj/20.8.1840. PubMed PMID: 11296218; PubMed Central PMCID: PMC125424.

73. Sheehan KB, Martin M, Lesser CF, Isberg RR, Newton IL. Identification and Characterization of a Candidate *Wolbachia pipientis* Type IV Effector That Interacts with the Actin Cytoskeleton. MBio. 2016;7(4). doi: 10.1128/mBio.00622-16. PubMed PMID: 27381293; PubMed Central PMCID: PMC4958246.

74. Sisko JL, Spaeth K, Kumar Y, Valdivia RH. Multifunctional analysis of *Chlamydia*-specific genes in a yeast expression system. Mol Microbiol. 2006;60(1):51–66. doi: 10.1111/j.1365-2958.2006.05074.x. PubMed PMID: 16556220.

75. Mittal E, Skowyra ML, Uwase G, Tinaztepe E, Mehra A, Koster S, et al. *Mycobacterium tuberculosis* Type VII Secretion System Effectors Differentially Impact the ESCRT Endomembrane Damage Response. mBio. 2018;9(6). Epub 20181127. doi: 10.1128/mBio.01765-18. PubMed PMID: 30482832; PubMed Central PMCID: PMC6282207.

76. Zhadina M, Bieniasz PD. Functional interchangeability of late domains, late domain cofactors and ubiquitin in viral budding. PLoS Pathog. 2010;6(10):e1001153. Epub 20101021. doi: 10.1371/journal.ppat.1001153. PubMed PMID: 20975941; PubMed Central PMCID: PMC2958808.

77. Demirov DG, Ono A, Orenstein JM, Freed EO. Overexpression of the N-terminal domain of TSG101 inhibits HIV-1 budding by blocking late domain function. Proc Natl Acad Sci U S A. 2002;99(2):955–60. doi: 10.1073/pnas.032511899. PubMed PMID: 11805336; PubMed Central PMCID: PMC117412.

78. Garrus JE, von Schwedler UK, Pornillos OW, Morham SG, Zavitz KH, Wang HE, et al. Tsg101 and the vacuolar protein sorting pathway are essential for HIV-1 budding. Cell. 2001;107(1):55–65. doi: 10.1016/s0092-8674(01)00506-2. PubMed PMID: 11595185.

79. VerPlank L, Bouamr F, LaGrassa TJ, Agresta B, Kikonyogo A, Leis J, et al. Tsg101, a homologue of ubiquitin-conjugating (E2) enzymes, binds the L domain in HIV type 1 Pr55(Gag). Proc Natl Acad Sci U S A. 2001;98(14):7724–9. Epub 20010626. doi: 10.1073/pnas.131059198. PubMed PMID: 11427703; PubMed Central PMCID: PMC35409.

80. Kikonyogo A, Bouamr F, Vana ML, Xiang Y, Aiyar A, Carter C, et al. Proteins related to the Nedd4 family of ubiquitin protein ligases interact with the L domain of Rous sarcoma virus and are required for gag budding from cells. Proc Natl Acad Sci U S A. 2001;98(20):11199–204. Epub 20010918. doi: 10.1073/pnas.201268998. PubMed PMID: 11562473; PubMed Central PMCID: PMC58707.

81. Martin-Serrano J, Eastman SW, Chung W, Bieniasz PD. HECT ubiquitin ligases link viral and cellular PPXY motifs to the vacuolar protein-sorting pathway. J Cell Biol. 2005;168(1):89–101. Epub 20041228. doi: 10.1083/jcb.200408155. PubMed PMID: 15623582; PubMed Central PMCID: PMC2171676.

82. Strack B, Calistri A, Accola MA, Palu G, Gottlinger HG. A role for ubiquitin ligase recruitment in retrovirus release. Proc Natl Acad Sci U S A. 2000;97(24):13063–8. doi: 10.1073/pnas.97.24.13063. PubMed PMID: 11087860; PubMed Central PMCID: PMC27178.

83. O’Connor HF, Lyon N, Leung JW, Agarwal P, Swaim CD, Miller KM, et al. Ubiquitin-Activated Interaction Traps (UBAITs) identify E3 ligase binding partners. EMBO Rep. 2015;16(12):1699–712. Epub 20151027. doi: 10.15252/embr.201540620. PubMed PMID: 26508657; PubMed Central PMCID: PMC4693525.

84. Ren J, Kee Y, Huibregtse JM, Piper RC. Hse1, a component of the yeast Hrs-STAM ubiquitin-sorting complex, associates with ubiquitin peptidases and a ligase to control sorting efficiency into multivesicular bodies. Mol Biol Cell. 2007;18(1):324–35. Epub 20061101. doi: 10.1091/mbc.e06-06-0557. PubMed PMID: 17079730; PubMed Central PMCID: PMC1751313.

85. Votteler J, Sundquist WI. Virus budding and the ESCRT pathway. Cell Host Microbe. 2013;14(3):232–41. doi: 10.1016/j.chom.2013.08.012. PubMed PMID: 24034610; PubMed Central PMCID: PMC3819203.

86. Rivera-Cuevas Y, Mayoral J, Di Cristina M, Lawrence AE, Olafsson EB, Patel RK, et al. *Toxoplasma gondii* exploits the host ESCRT machinery for parasite uptake of host cytosolic proteins. PLoS Pathog. 2021;17(12):e1010138. Epub 20211213. doi: 10.1371/journal.ppat.1010138. PubMed PMID: 34898650; PubMed Central PMCID: PMC8700025.

87. Guerin A, Corrales RM, Parker ML, Lamarque MH, Jacot D, El Hajj H, et al. Efficient invasion by *Toxoplasma* depends on the subversion of host protein networks. Nat Microbiol. 2017;2(10):1358–66. Epub 20170828. doi: 10.1038/s41564-017-0018-1. PubMed PMID: 28848228.

88. Mayoral J, Guevara RB, Rivera-Cuevas Y, Tu V, Tomita T, Romano JD, et al. Dense Granule Protein GRA64 Interacts with Host Cell ESCRT Proteins during *Toxoplasma gondii* Infection. mBio. 2022;13(4):e0144222. Epub 20220622. doi: 10.1128/mbio.01442-22. PubMed PMID: 35730903; PubMed Central PMCID: PMC9426488.

89. Grobler Y, Yun CY, Kahler DJ, Bergman CM, Lee H, Oliver B, et al. Whole genome screen reveals a novel relationship between *Wolbachia* levels and *Drosophila* host translation. PLoS Pathog. 2018;14(11):e1007445. Epub 20181113. doi: 10.1371/journal.ppat.1007445. PubMed PMID: 30422992; PubMed Central PMCID: PMC6258568.

90. Voronin D, Cook DA, Steven A, Taylor MJ. Autophagy regulates *Wolbachia* populations across diverse symbiotic associations. Proc Natl Acad Sci U S A. 2012;109(25):E1638–46. doi: 10.1073/pnas.1203519109. PubMed PMID: 22645363; PubMed Central PMCID: PMC3382551.

91. Takahashi Y, He H, Tang Z, Hattori T, Liu Y, Young MM, et al. An autophagy assay reveals the ESCRT-III component CHMP2A as a regulator of phagophore closure. Nat Commun. 2018;9(1):2855. Epub 20180720. doi: 10.1038/s41467-018-05254-w. PubMed PMID: 30030437; PubMed Central PMCID: PMC6054611.

92. Zhen Y, Spangenberg H, Munson MJ, Brech A, Schink KO, Tan KW, et al. ESCRT-mediated phagophore sealing during mitophagy. Autophagy. 2020;16(5):826–41. Epub 20190801. doi: 10.1080/15548627.2019.1639301. PubMed PMID: 31366282; PubMed Central PMCID: PMC7158923.

93. Schafer JA, Schessner JP, Bircham PW, Tsuji T, Funaya C, Pajonk O, et al. ESCRT machinery mediates selective microautophagy of endoplasmic reticulum in yeast. EMBO J. 2020;39(2):e102586. Epub 20191205. doi: 10.15252/embj.2019102586. PubMed PMID: 31802527; PubMed Central PMCID: PMC6960443.

94. Quek S, Cook DAN, Wu Y, Marriott AE, Steven A, Johnston KL, et al. *Wolbachia* depletion blocks transmission of lymphatic filariasis by preventing chitinase-dependent parasite exsheathment. Proc Natl Acad Sci U S A. 2022;119(15):e2120003119. Epub 20220404. doi: 10.1073/pnas.2120003119. PubMed PMID: 35377795; PubMed Central PMCID: PMC9169722.

95. Cascales E, Bernadac A, Gavioli M, Lazzaroni JC, Lloubes R. Pal lipoprotein of *Escherichia coli* plays a major role in outer membrane integrity. J Bacteriol. 2002;184(3):754–9. doi: 10.1128/JB.184.3.754-759.2002. PubMed PMID: 11790745; PubMed Central PMCID: PMC139529.

96. Szczepaniak J, Press C, Kleanthous C. The multifarious roles of Tol-Pal in Gram-negative bacteria. FEMS Microbiol Rev. 2020;44(4):490–506. doi: 10.1093/femsre/fuaa018. PubMed PMID: 32472934; PubMed Central PMCID: PMC7391070.

97. Tan WB, Chng SS. Genetic interaction mapping highlights key roles of the Tol-Pal complex. Mol Microbiol. 2022;117(4):921–36. Epub 20220221. doi: 10.1111/mmi.14882. PubMed PMID: 35066953.

98. Yakhnina AA, Bernhardt TG. The Tol-Pal system is required for peptidoglycan-cleaving enzymes to complete bacterial cell division. Proc Natl Acad Sci U S A. 2020;117(12):6777–83. Epub 20200309. doi: 10.1073/pnas.1919267117. PubMed PMID: 32152098; PubMed Central PMCID: PMC7104345.

99. Foster J, Ganatra M, Kamal I, Ware J, Makarova K, Ivanova N, et al. The *Wolbachia* genome of *Brugia malayi*: endosymbiont evolution within a human pathogenic nematode. PLoS Biol. 2005;3(4):e121. doi: 10.1371/journal.pbio.0030121. PubMed PMID: 15780005; PubMed Central PMCID: PMC1069646.

100. Michel LV, Shaw J, MacPherson V, Barnard D, Bettinger J, D’Arcy B, et al. Dual orientation of the outer membrane lipoprotein Pal in *Escherichia coli*. Microbiology. 2015;161(6):1251–9. Epub 2015/03/27. doi: 10.1099/mic.0.000084. PubMed PMID: 25808171; PubMed Central PMCID: PMC4635515.

101. Voronin D, Guimaraes AF, Molyneux GR, Johnston KL, Ford L, Taylor MJ. *Wolbachia* lipoproteins: abundance, localisation and serology of *Wolbachia* peptidoglycan associated lipoprotein and the Type IV Secretion System component, VirB6 from *Brugia malayi* and *Aedes albopictus*. Parasit Vectors. 2014;7:462. doi: 10.1186/s13071-014-0462-1. PubMed PMID: 25287420; PubMed Central PMCID: PMC4197220.

102. Foray V, Perez-Jimenez MM, Fattouh N, Landmann F. *Wolbachia* Control Stem Cell Behavior and Stimulate Germline Proliferation in Filarial Nematodes. Dev Cell. 2018;45(2):198–211 e3. Epub 2018/04/25. doi: 10.1016/j.devcel.2018.03.017. PubMed PMID: 29689195.

103. Carlton JG, Martin-Serrano J. Parallels between cytokinesis and retroviral budding: a role for the ESCRT machinery. Science. 2007;316(5833):1908–12. Epub 20070607. doi: 10.1126/science.1143422. PubMed PMID: 17556548.

104. Morita E, Sandrin V, Chung HY, Morham SG, Gygi SP, Rodesch CK, et al. Human ESCRT and ALIX proteins interact with proteins of the midbody and function in cytokinesis. EMBO J. 2007;26(19):4215–27. Epub 20070913. doi: 10.1038/sj.emboj.7601850. PubMed PMID: 17853893; PubMed Central PMCID: PMC2230844.

105. Gao CY, Pinkham JL. Tightly regulated, beta-estradiol dose-dependent expression system for yeast. Biotechniques. 2000;29(6):1226–31. PubMed PMID: 11126125.

106. Mumberg D, Muller R, Funk M. Yeast vectors for the controlled expression of heterologous proteins in different genetic backgrounds. Gene. 1995;156(1):119–22. doi: 10.1016/0378-1119(95)00037-7. PubMed PMID: 7737504.

107. Gietz RD, Schiestl RH. High-efficiency yeast transformation using the LiAc/SS carrier DNA/PEG method. Nat Protoc. 2007;2(1):31–4. doi: 10.1038/nprot.2007.13. PubMed PMID: 17401334.

108. Macreadie IG, Horaitis O, Verkuylen AJ, Savin KW. Improved shuttle vectors for cloning and high-level Cu(2+)-mediated expression of foreign genes in yeast. Gene. 1991;104(1):107–11. doi: 10.1016/0378-1119(91)90474-p. PubMed PMID: 1916270.

109. Altschul SF, Gish W, Miller W, Myers EW, Lipman DJ. Basic local alignment search tool. J Mol Biol. 1990;215(3):403–10. Epub 1990/10/05. doi: 10.1016/S0022-2836(05)80360-2. PubMed PMID: 2231712.

110. Sternberg PW, Van Auken K, Wang Q, Wright A, Yook K, Zarowiecki M, et al. WormBase 2024: status and transitioning to Alliance infrastructure. Genetics. 2024;227(1). doi: 10.1093/genetics/iyae050. PubMed PMID: 38573366; PubMed Central PMCID: PMC11075546.

111. Goldstein AL, McCusker JH. Three new dominant drug resistance cassettes for gene disruption in *Saccharomyces cerevisiae*. Yeast. 1999;15(14):1541–53. doi: 10.1002/(SICI)1097-0061(199910)15:14<1541::AID-YEA476>3.0.CO;2-K. PubMed PMID: 10514571.

112. von der Haar T. Optimized protein extraction for quantitative proteomics of yeasts. PLoS One. 2007;2(10):e1078. Epub 20071024. doi: 10.1371/journal.pone.0001078. PubMed PMID: 17957260; PubMed Central PMCID: PMC2031916.

113. Schindelin J, Arganda-Carreras I, Frise E, Kaynig V, Longair M, Pietzsch T, et al. Fiji: an open-source platform for biological-image analysis. Nat Methods. 2012;9(7):676–82. Epub 2012/06/30. doi: 10.1038/nmeth.2019. PubMed PMID: 22743772; PubMed Central PMCID: PMC3855844.

114. Schneider CA, Rasband WS, Eliceiri KW. NIH Image to ImageJ: 25 years of image analysis. Nat Methods. 2012;9(7):671–5. Epub 2012/08/30. PubMed PMID: 22930834; PubMed Central PMCID: PMC5554542.

115. Giddings TH. Freeze-substitution protocols for improved visualization of membranes in high-pressure frozen samples. J Microsc. 2003;212(Pt 1):53–61. doi: 10.1046/j.1365-2818.2003.01228.x. PubMed PMID: 14516362.

116. Mastronarde DN. Dual-axis tomography: an approach with alignment methods that preserve resolution. J Struct Biol. 1997;120(3):343–52. doi: 10.1006/jsbi.1997.3919. PubMed PMID: 9441937.

117. Mastronarde DN. Automated electron microscope tomography using robust prediction of specimen movements. J Struct Biol. 2005;152(1):36–51. doi: 10.1016/j.jsb.2005.07.007. PubMed PMID: 16182563.

118. Kremer JR, Mastronarde DN, McIntosh JR. Computer visualization of three-dimensional image data using IMOD. J Struct Biol. 1996;116(1):71–6. doi: 10.1006/jsbi.1996.0013. PubMed PMID: 8742726.

